# Deconvolving nuclesome binding energy from experimental errors

**DOI:** 10.1101/2020.02.20.957878

**Authors:** Mark Heron, Johannes Soeding

## Abstract

Eukaryotic genomes are compacted into nucleosomes, 147-bp of DNA wrapped around histone proteins. Nucleosomes can hinder transcription factors and other DNA-binding proteins from accessing the genome. This competition at promoters and enhancers regulates gene expression. Therefore, a quantitative understanding of gene regulation requires the quantitative prediction of nucleosome binding affinities. However, little is known for certain about the sequence preference of nucleosomes.

Here we develop an integrated model of nucleosome binding and genome-wide measurements thereof. Our model learns similar nucleosome sequence preferences from MNase-Seq and CC-Seq datasets.

We find that modelling the positional uncertainty of MNase-Seq deconvolves the commonly described smooth 10-bp-periodic sequence preference into a position-specific pattern more closely resembling the pattern obtained from high-resolution CC-Seq data. By analysing the CC-Seq data we reveal the strong preference of A/T at +/− 3 bp from the dyad as an experimental bias. Our integrated model can separate this bias of CC-Seq from the true nucleosome binding preference.

Our results show that nucleosomes have position-specific sequence preferences, which probably play an important role in their competition with transcription factors. Furthermore, our comparison of diverse datasets shows that the experimental biases have a similar strength as the signal of nucleosome-positioning measurements. Validating nucleosome models on experiments with similar biases overestimates their prediction quality of the true nucleosome binding.

There are still many open questions about the sequence preference of nucleosomes and our approach will need to be extended to answer them. Only integrated models that combine the thermodynamics of nucleosome binding with experimental errors can deconvolve the two and learn the true preferences of nucleosomes.

## 2 Introduction

Eukaryotic genomes are packed into chromatin. The basic unit of chromatin is a nucleosome – 147 bps of DNA wrapped around a histone octamer. This wrapping strongly affects the accessibility of the nucleosomal DNA. Most transcription factors primarily bind accessible regions of the genome, which are unoccupied by nucleosomes. The competition between the binding of nucleosomes and transcription factors at promoters regulates gene expression (Raveh-Sadka et al. 2012). To decode the complex interactions at promoters and predict transcription-factor occupancy and the resulting gene expression from sequence alone, we need to understand the sequence features nucleosomes preferentially bind to (Segal and Widom 2009; Iyer 2012).

The primary determinants of *in vitro* nucleosome formation are the bendability of the DNA sequence and steric hindrance between neighboring nucleosomes. The first genome-wide measurements of nucleosomes led to disagreements about the existence of a ‘nucleosome positioning code’, i.e. whether DNA sequence primarily determines the nucleosome positioning (Segal et al. 2006; Kaplan et al. 2009; Tillo and Hughes 2009; Noam Kaplan, Moore, et al. 2010) or whether sequence is but one of various factors (Zhang et al. 2009; Stein, Takasuka, and Collings 2010; Locke et al. 2010; Zhang et al. 2010; Chung et al. 2010). Later reviews unified the different interpretations by distinguishing between local rotational positioning (conditional positioning) and genome-wide positioning (absolute positioning – which we use throughout the paper), and acknowledging experimental limitations (Noam Kaplan, Hughes, et al. 2010; Iyer 2012; Struhl and Segal 2013). On the one hand, genome-wide occupancy can be predicted well by sequence, but the correlations are inflated due to biases of MNase-Seq. On the other hand, rotational positioning cannot be predicted well at base-pair resolution.

*In vivo* the positioning of nucleosomes results from interactions between nucleosome remodelers, competition with transcription factors and other DNA-binding proteins, Pol II elongation, steric hindrance between nucleosomes, and their DNA sequence binding preferences. Nucleosomes are believed to prefer higher G+C content regions, avoid homo-polymeric sequences (both poly(dA:dT) and poly(dC:dG)), and their rotational position relies on a 10-bp-periodic pattern of SS to WW enrichment (W is A or T and S is C or G). These sequence preferences are frequently overridden – primarily at promoters and enhancers – by nucleosome remodelers and competition with transcription factors.

The most common technique to measure genome-wide nucleosome binding is MNase-Seq. The chromatin is digested with Micrococcal nuclease (MNase), and fragments of about the length of a nucleosome (147 bps) are isolated and sequenced. Other experimental techniques measure nucleosome binding genome-wide, e.g. MPE-Seq (Ishii, Kadonaga, and Ren 2015), RED-Seq (Chen et al. 2014), NOMe-Seq (Kelly et al. 2012); nucleosome binding at individual loci, e.g. by electron microscopy (Brown et al. 2013), by DNA methylation (Small et al. 2014); or nucleosome binding to short DNA fragments *in vitro*, e.g. BunDLE-Seq (Levo et al. 2015). CC-Seq (Chemical Cleavage with sequencing, also known as HC-Seq or Chemical Map) stands out amongst the genome-wide *in vivo* measurements due to its high positional resolution (Brogaard et al. 2012). A copper-chelating label is covalently bound to the genetically modified H4S47C. By adding copper and hydrogen, hydroxyl radicals form in proximity to the nucleosome dyad and cleave the DNA backbone at specific positions. Fragments spanning between two cleavage sites – i.e. nucleosome dyads – are isolated and sequenced. CC-Seq has not yet been studied extensively: it has a high positional resolution, but possible biases have not been analysed much.

MNase-Seq has been studied extensively, revealing several limitations: a limited positional resolution (Noam Kaplan, Hughes, et al. 2010), a dependency on the degree of digestion (Weiner et al. 2010; Rizzo, Bard, and Buck 2012), and an unsettling, high correlation to nucleosome-free MNase-Seq experiments (Locke et al. 2010; Chung et al. 2010). The low positional resolution limits the analysis of rotational sequence preferences to the 10-bp-periodic SS/WW enrichments (Heijden et al. 2012; Struhl and Segal 2013), while the analysis of CC-Seq data later revealed non-periodic preferences (Brogaard et al. 2012). The dependency on the degree of digestion means that a single MNase-Seq experiment only captures a fraction of all nucleosomes based on their stability. These limitations of MNase-Seq have often been found unimportant for the interpretation of individual results (Noam Kaplan, Moore, et al. 2010; Zentner and Henikoff 2012), but when trying to understand and predict nucleosome positioning in a quantitative fashion such limitations need to be taken into account (Chung et al. 2010).

The high correlations of MNase-Seq measurements to nucleosome-free control experiments must stem from experimental biases. We re-analysed published datasets in a critical and comprehensive way to investigate such experimental biases. Correlations between the datasets reveal strong biases that dominate all genome-wide nucleosome measurements. These biases manifest themselves in unrealistic occupancy distributions given the standard assumption that the measured nucleosome binding is proportional to the probability of nucleosomes binding *in vivo*. All of this can cause problems when modeling and predicting nucleosome binding.

A variety of computational methods exist that predict different aspects of nucleosome binding (Liu et al. 2014; Teif 2015). Some methods predict absolute nucleosome-positioning scores for whole genomes. Most methods consist of two steps: learning nucleosome sequence preferences from measured nucleosome positions, and predicting the occupancies for a query sequence using a thermodynamic model that includes steric hindrance. The first such predictor obtained the sequence preferences by deriving the position-specific enrichment of dinucleotides from their frequencies in aligned binding sites (Segal et al. 2006). Two noteworthy extensions are: accounting for competing local nucleosomes when extracting the sequence preferences in the first step (Locke et al. 2010), and adding competition with transcription factors to the thermodynamic model (Wasson and Hartemink 2009).

Published nucleosome-position predictors are good at reproducing genome-wide MNase-Seq measurements at low positional resolution (Kaplan et al. 2009; Tillo and Hughes 2009). However, it is unclear how close these predictions are to the actual nucleosome positioning, due to the unknown experimental biases. Also, the positional resolution of the predictions is lower than that of the rotational positioning (Locke et al. 2010). This restricts the usefulness of these nucleosome-position predictors in decoding the complex interactions involved in nucleosome formation *in vivo*.

Here we present a novel integrated approach to predicting nucleosome positioning. Our model distinguishes between nucleosome binding events and measurements of these events, which allows us to deconvolve the two. We integrate the thermodynamic model describing nucleosome binding into a probabilistic model that adds the experimental biases and positional uncertainty of the measurements. By maximizing a likelihood of observing the experimental data, we simultaneously learn the nucleosome binding preference, experimental bias, and positional uncertainty in our integrated model.

Thanks to the deconvolution approach, our method can learn a high-resolution nucleosome binding energy model from low-resolution MNase-Seq data. Compared to previous predictors it has the highest correlation to CC-Seq data at base-pair resolution. We show that the CC-Seq experiment has a sequence bias and our method separates this bias from the nucleosome binding energies. With these improvements the two nucleosome binding energy models derived from the distinct experiments converge. Both have highly position-specific sequence preferences that correlate strongly with each other. This confirms that the smoothness of the commonly described 10-bp-periodic SS and WW sequence preference stems from the low positional resolution of MNase-Seq.

Our approach to integrate the thermodynamics into a probabilistic model of the measurements provides a blueprint to separate the relevant signal from experimental errors when extracting nucleosome binding energies. We demonstrate the improvements with our computational method that can correct for the lower positional resolution of MNase-Seq and the sequence bias of CC-Seq data. The higher similarity of our nucleosome binding energy models suggests they are closer to the real binding energies than uncorrected models.

## 3 Methods

In contrast to previous approaches our probabilistic model does not assume that the nucleosome binding positions are given directly by the measurements. Rather, the measured distribution depends on the true binding positions, the positional error, and the sequence bias. Figure 1A shows a schematic of our probabilistic method with the nucleosome-positioning model colored in purple and the data model colored in green. The likelihood of our model for the experimental measurements is:

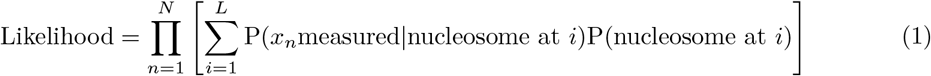

P(nucleosome at *i*) is the probability of a nucleosome binding at position *i* – the nucleosome-positioning model. P(*x*_*n*_measured nucleosome at *i*) is the the probability of a measurement observed at position *x*_*n*_, given a nucleosome at *i* – the experimental error model. *L* is the sequence length and *N* is the amount of measurements.

**Figure 1:**
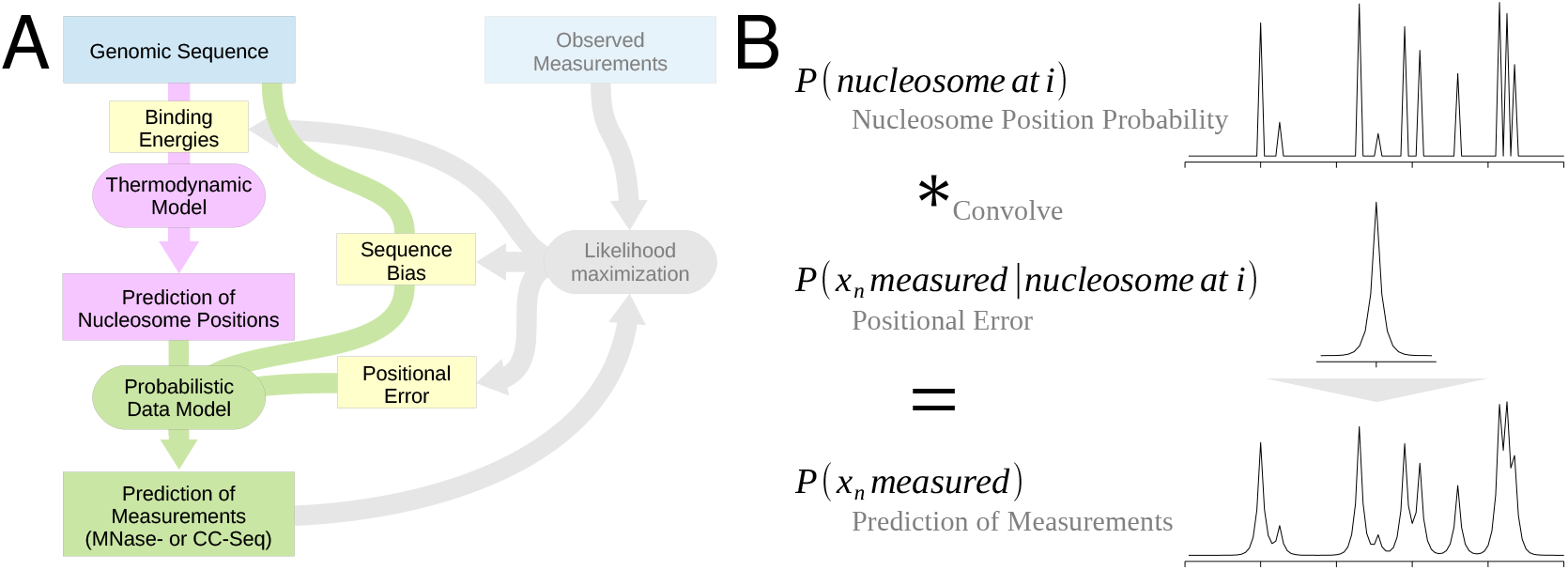
Our model distinguishes between a nucleosome position and its experimental measurement. (**A**) Our probabilistic model consists of a thermodynamic part (purple) that models the nucleosome positioning, and a probabilistic-data part (green) that can model experimental sequence biases (CC-Seq) and positional errors (primarily MNase-Seq) of the measurements. All parameters of our model (yellow) are optimized by maximizing the likelihood of the observed measurements (gray). (**B**) The predicted distribution of measurements results from the convolution of the predicted nucleosome binding sites with the positional error function.

The experimental error model P(*x*_*n*_measured nucleosome at *i*) can describe a position error function, a sequence dependent experimental bias, or both. We developed two position error functions: a Laplace distribution representing positional noise and a position specific pattern, which can learn CC-Seq’s cut pattern. Equation 2 shows the basic positional noise and Figure 1B visualizes the effect it has. The position specific convolution pattern and the sequence dependent bias term used in the CC-Seq model are described in the Supplemental Methods.

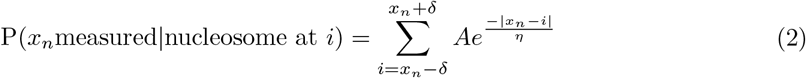

*η* is the scaling factor or width of the distribution, *A* is the normalization term and *δ* the fixed maximum distance modeled between a nucleosome and a measurement (set to 20 bps based on the irrelevance of larger distances).

The nucleosome-positioning model P(nucleosome at *i*) is a thermodynamic model that includes steric hindrance between neighboring nucleosomes. The model describes the probability of a nucleosome dyad occurring at position *i* compared to other genomic positions.

Our method optimizes the nucleosome binding energies, which are part of the thermodynamic model, in the context of the whole probabilistic model, which includes the experimental errors. To optimize the model parameters the method maximizes the likelihood of observing the measured nucleosome positions (*x*_*n*_). We use a gradient ascent for the optimization, because the likelihood’s derivatives can be calculated for the parameters (see Supplemental Methods).

Our model represent the sequence-specific binding energy with a Markov chain of 1st-order, i.e. the probabilities depend on one previous position. To represent the dyad symmetric nature of nucleosome binding our method uses conditional probabilities that depend on the positions towards the dyad. The energy terms *ε* that encode the Markov chain represent the logarithm of the conditional probabilities.

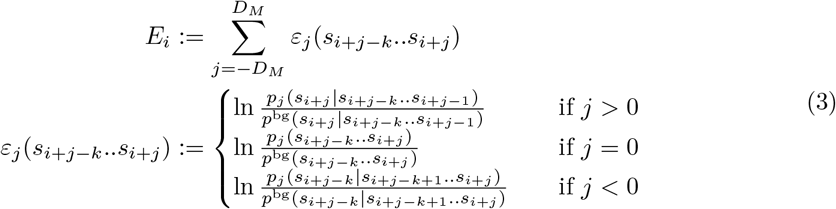

*k* is the order of the Markov chain and *D*_*M*_ is the half size of the energy model. The method initializes the parameters *ε* based on Equation 3 by estimating the probabilities *p* and genomic background probabilities *p*^bg^ from nucleotide frequencies. Assuming that the data has position-specific resolution and the nucleosome binding preference only depends on the wrapped sequence, the nucleosome binding energy can be derived from the binding frequencies following Bolzmann’s law. The parameters 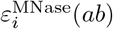 of the 1st-order nucleosome binding energy model are initialized as the log ratio between the conditional frequency of dinucleotide *ab* at position *i* and the genomic background frequency (see Supplemental Methods 1.2.1). The case separation conserves the dyad symmetry of the energy-model parameters.

In the model *µ* represents the sequence-unspecific binding energy. It contains the general binding preference or chemical potential of a nucleosome, and the nucleosome concentration. Because the two cannot be separated without measuring either in an independent experiment, the model represents both with a single parameter.

## 4 Results

### 4.1 Strong biases dominate genome-wide nucleosome occupancy measurements

We systematically compared nucleosome datasets of several publications with Pearson’s correlation coefficient between the derived genomic occupancies, as proposed by Noam Kaplan, Hughes, et al. (2010). Nucleosome occupancy describes the probability with which a nucleosome covers a genomic position (see Noam Kaplan, Hughes, et al. (2010) for a formal definition). We selected datasets that use distinct measurement methods to analyze a broad spectrum of experiments (Brogaard et al. 2012; Kaplan et al. 2009; Field et al. 2008; Mavrich et al. 2008; Lee et al. 2007; Fan et al. 2010; Zhang et al. 2009). We added three MNase-Seq datasets with a knockdown or heat shock treatment, which are expected to affect genome-wide nucleosome positioning, in order to compare the experimental bias and signal strengths (Shivaswamy et al. 2008; Gossett and Lieb 2012; Celona et al. 2011). A sample of the compared genomic occupancy tracks is given in Figure 2B.

**Figure 2:**
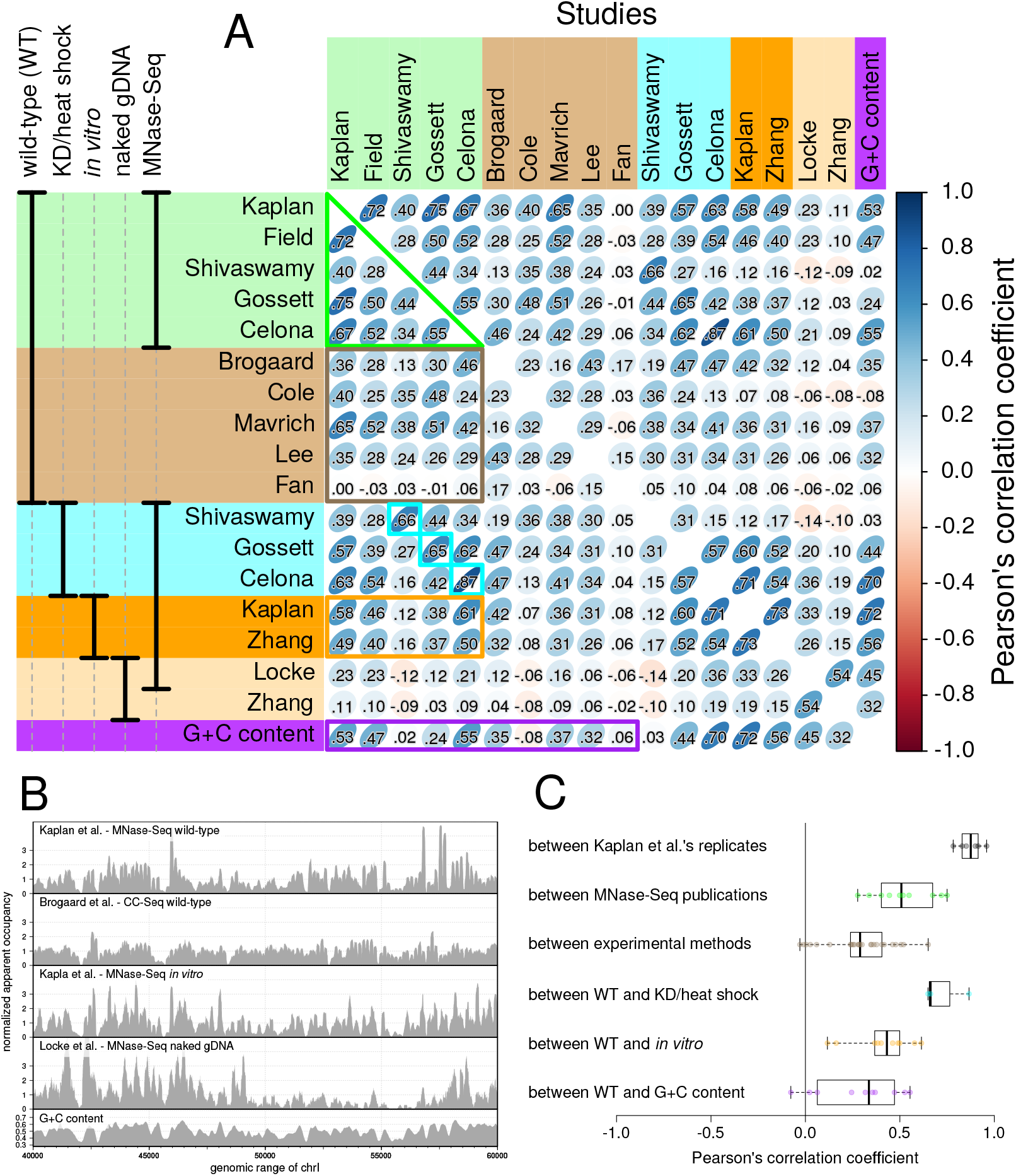
Correlations between experimental nucleosome data reveal strong biases. (**A**) Matrix of Pearson’s correlation coefficients between occupancies derived from nucleosome measurements, control experiments and G+C content for *S.cerevisia*. The annotation on the left describes shared features of the used experiments. (**B**) Individual occupancy tracks for a subset of the measurements and G+C content. (**C**) Boxplots of grouped comparisons outlined by the same color in (A).

The correlation coefficients between genomic nucleosome occupancies measured by MNase-Seq of untreated wild-type cells in different studies are generally low (median of 0.51; green triangle in Figure 2A) given that they effectively represent biological replicate measurements (Supplement Table S1). The replicates from the Kaplan et al. (2009) publication have higher correlations with each other (median of 0.88; Figure 2C). This suggests that the low correlations are not due to experimental noise but due to technical details that strongly influence measurement biases. The correlations between measurements of MNase-Seq and other experimental protocols (median of 0.29; brown rectangle in Figure 2A) are even lower than those between the MNase-Seq measurements, and lower than correlations between wild-type and *in vitro* MNase-Seq measurements (median of 0.46, orange frame in Figure 2A).

Correlations between datasets of untreated wild-type and heat shock or knock-down cells from the same publication are among the highest (compare cyan to the other values in Figure 2A,C), even though the knocked-down chromosomal proteins (H3 and nhp6) take part in nucleosome formation (Gossett and Lieb 2012; Celona et al. 2011). Batch effects, which lead to increased correlations in datasets produced in one batch or from one laboratory, are known from gene expression measurements (Leek et al. 2010). These results show that they affect MNase-Seq measurements as well. The batch effects are also partially responsible for the stronger correlations between the replicate measurements than between the MNase-Seq measurements from different publications.

The correlation of wild-type nucleosome measurements to G+C content varies strongly (purple in Figure 2A,C), and the measurements therefore disagree on the importance of the most basic sequence feature. Three datasets have reasonable correlations (*r* = 0.53, 0.47, and 0.55) suggesting the G+C content is a major determining feature of nucleosome positioning. However, three datasets have barely any or negative correlations (*r* = 0.02, −0.08, and 0.06), which would suggest that nucleosomes are positioned independent of G+C content.

Our analysis shows that the experimental biases are of the same order of magnitude or larger than the observed difference between wild-type and knock-down, heat shock or *in vitro* measurements. Any information gained from the measurements without separating out the biases contains at least as much bias as signal. One illustration of this issue is the strong variation of the measurement’s correlation to G+C content, which ranges from *r* = *−*0.08 to 0.55.

### 4.2 Strong biases in measured genomic nucleosome binding manifest themselves in unrealistic occupancy distributions

Segal et al. (2006) estimated the average nucleosome occupancy in the range of 75-90% and we estimate it in the range of 68-83% (see Supplemental Information). Such average occupancies set a hard upper limit for fully occupied DNA at 1.2-1.5 times the average. To better understand what occupancy distribution is expected, we computed distributions predicted by our model, which includes steric exclusion and nucleosome sequence preferences. By setting the unspecific binding energy (*µ*) the average occupancy is influenced. We chose a realistic average occupancy of 78% (*µ* = 0), and to match the observed distribution better we also chose an unrealistically-low average occupancy of 41% (*µ* = 4) (Figure 3A,B). The occupancy distribution stays below 1.3 times the average for our realistic prediction.

**Figure 3:**
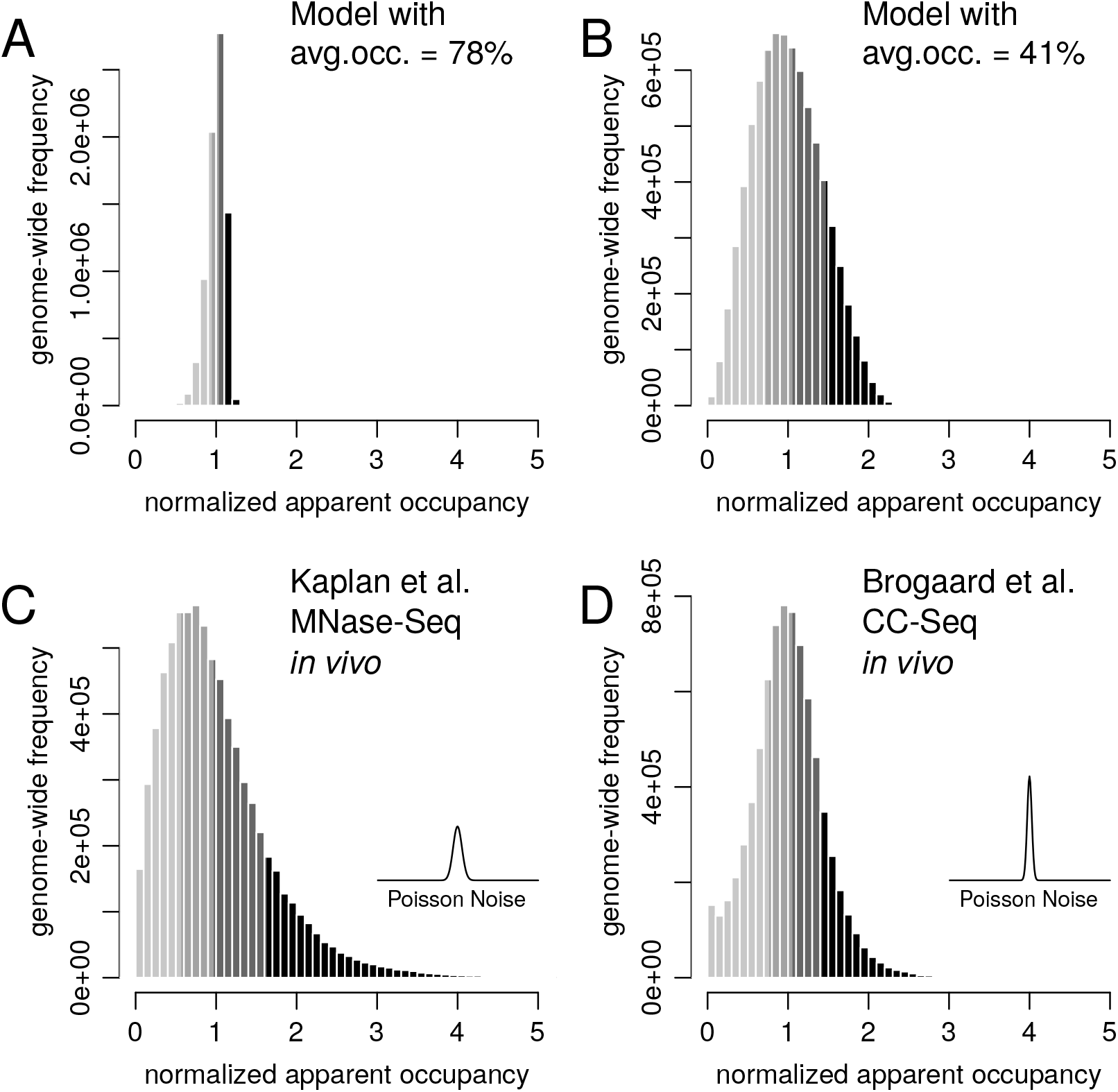
Nucleosome occupancies derived from MNase-Seq and CC-Seq have an unrealistic distributions. Histograms of nucleosome occupancies derived from our model predictions (**A**, **B**) and measured binding events (**C**, **D**). Assuming the measured nucleosome binding is proportional to the true nucleosome binding probability leads to an unrealistically-low average nucleosome occupancy. The occupancies are normalized to a mean of 1 and the four shades depict the quartiles. (A) and (B) show predictions of our method without added Poisson noise. We set the unspecific binding-energy parameter of our model to produce a realistic 78% (A) and an unrealistically-low 41% (B) occupancy. The inserts in (C) and (D) depict the expected Poisson noise at a nor. app. occ. of 1, with the same scale as the x-axis.

To investigate the effects of possible biases, we checked the occupancy distributions of the genome-wide measurements normalized to a mean of 1 (Figure 3C,D and Supplemental Figure S1). In interpreting nucleosome measurements, it is usually implicitly assumed that the density of the measured nucleosome positions is proportional to the actual nucleosome binding frequency over the genome. Therefore, the occupancy would be proportional to a 147-bp convolution of the measured density. The distributions of all measurements are at least as wide as the distribution of the unrealistically-low occupancy prediction and in most cases even the 75% quantile is above the hard upper limit of 1.5. These heavy tails cannot be explained by sampling noise (line inserts in Figure 3C,D).

There are two possible explanations for the discrepancy between the expected and observed distributions: either the published and our genomic occupancy estimates are off by more than a factor of two, or the signals measured by MNase-Seq, CC-Seq, etc. are biased measurements of nucleosome binding. The latter explanation is more probable given that our analysis above showed that the signal and bias of the measurements are of the same order of magnitude. It has been suggested before that read frequencies of MNase-Seq are not quantitative representations of nucleosome binding frequencies (Zhang et al. 2009). Due to these shortcomings, we decided to focus on obtaining a positionally-high-resolution nucleosome energy model instead of reproducing measured occupancies.

### 4.3 A high resolution nucleosome energy model from MNase-Seq data

When analyzing experimental data of nucleosome positioning, a common implicit assumption is that the measurements have base-pair resolution. In contrast, our approach explicitly models the probability of observing a certain data point given the nucleosome positioning. The first part of our method computes the probability of each genome position to be covered by a nuclesome dyad using a thermodynamic model (orange part in Figure 1A). The method further computes the probability of measuring a data point at a genomic position – here MNase-Seq reads (green part in Figure 1A). These MNase-Seq model probabilities contain the positional uncertainty of MNase-Seq and are therefore equivalent to the predictions of other methods that are trained on MNase-Seq data and do not distinguish between a nucleosome position and its measurement (e.g. Kaplan et al.’s method). The distinction between the positioning of nucleosomes and their experimental measurement allows us to obtain a high-resolution energy model from low-resolution MNase-Seq data. Our method iteratively learns both the nucleosome binding energy model and the positional-uncertainty model by maximizing a log-likelihood (Methods and Supplemental Methods).

To compare our method to Kaplan et al.’s method (Kaplan et al. 2009) we trained our model parameters on their *in vitro* MNase-Seq dataset, which they also used to train their model. We compared the methods by computing Pearson’s correlation coefficients between the predicted probabilities and several test datasets (Figure 4 and Supplemental Figure S2). We correlated dyad frequencies (i.e. base-pair resolution) – not occupancies (i.e. 147-bp smoothed) – to highlight the resolution of the predictions. Our method has higher correlations than Kaplan et al.’s method to most datasets and comparable correlations otherwise. NuPoP (Xi et al. 2010), another nucleosome prediction method we tested, has very low correlations in our comparison (Supplemental Figure S2).

**Figure 4:**
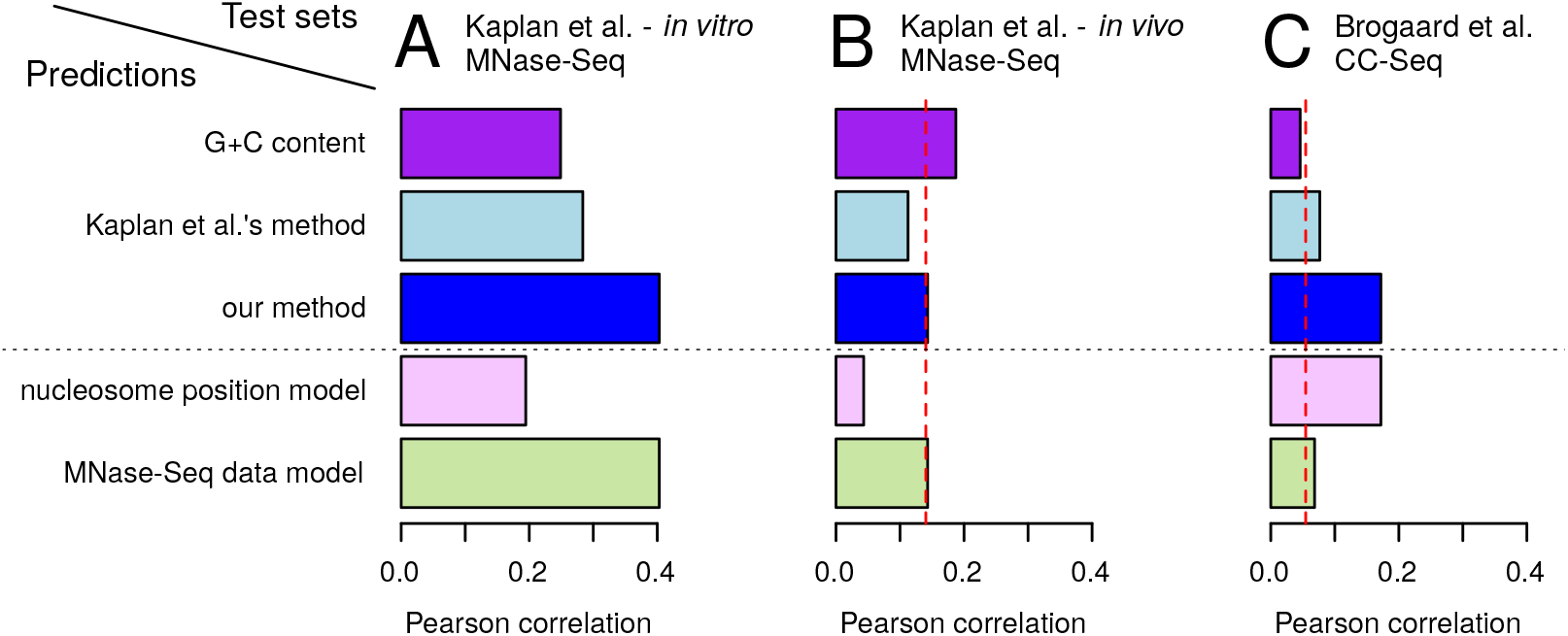
Our method performs well at predicting nucleosome positioning at basepair resolution. Pearson’s correlation coefficients between prediction methods (rows) and experimental measurements (columns **A**, **B**, **C**). Kaplan et al.‘s method and our method were trained on the *in vitro* dataset of Kaplan et al.. The correlation of this dataset to the other two is shown by the dashed red lines. ’G+C content’ uses the G+C content of the surrounding 147-bp window as a predictor. ‘Our method’ (blue) is the relevant prediction of our model for the given test set. Below the dashed line we show both predictions of our model: the MNase-Seq data prediction (green) and nucleosome-position predictions (pink).

Both the method of Kaplan et al. and ours perform best on their *in vitro* dataset (Figure 4A). This is expected given that the energy models of both methods are based on this dataset. The correlations of the two methods predictions to MNase-Seq test sets follows the trend of the prediction quality of G+C content (Figure 4A,B and Supplemental Figure S2). This suggests that even at single-base-pair resolution the methods’ performances rely strongly on the average G+C content of the nucleosome footprint.

The low-positional resolution of MNase-Seq lead to higher correlations to predictions that have a similarly low resolution. This becomes apparent when comparing the two probabilities of our model trained on MNase-Seq data with the MNase-Seq test sets (Figure 4A,B below the dotted line): the correlations of our MNase-Seq-data model probabilities are more than double the corresponding correlations of our nucleosome-position probabilities. In other words, adding the positional uncertainty – the only difference between the two probabilities – improves the correlation coefficients while decreasing the prediction’s resolution.

Data with strong biases can also make benchmarks of prediction methods misleadingly optimistic. This issue occurs if the training data, on which a method’s parameters are optimized, contains the same strong bias as the test data that is used to estimate the method’s performance. The prediction scores are inflated by predicting the biases shared by the training and test data. To mitigate this problem the training and test datasets should be measurements from two methods that have different biases. For this reason, we used an *in vivo* CC-Seq dataset (Brogaard et al. 2012) as a test set whose experimental protocol is very different to that of the MNase-Seq training set (Figure 4C). While CC-Seq and MNase-Seq both have a sequencing step, which can produce sequence-dependent biases (Harismendy et al. 2009), CC-Seq is otherwise distinct from MNase-Seq: instead of MNase digesting linker DNA the nucleosomal DNA is cleaved close to the dyad. Due to CC-Seq’s high resolution in comparison to MNase-Seq, our nucleosome-position prediction doubles the correlation coefficients of the predictions of Kaplan et al. and our MNase-Seq-data model, which both contain the positional uncertainty of MNase-Seq. This confirms that our approach improves the learned nucleosome binding preferences by explicitly modeling the positional uncertainty.

### 4.4 CC-Seq has a strand-specific bias close to the dyad position

While the CC-Seq dataset published by Brogaard et al. has a higher positional resolution than MNase-Seq data, we showed that neither have genome-wide distributions consistent with unbiased relative nucleosome occupancies (Figure 3). In comparison to nucleosomes measured with MNase-Seq, CC-Seq produces an A and T enrichment at the −3 and +3 positions from the nucleosome dyad, respectively. The authors originally conceded that this could be a bias (Brogaard et al. 2012), but later claimed it validated an A/T enrichment seen in a previous study (Xi et al. 2014). Cole et al. (2015) provided several possible explanations for this A/T enrichment based on biases.

We show here that the enrichments are indeed caused by an experimental bias. In CC-Seq, the genome-wide fragment end counts are deconvolved based on the distribution of cuts around the nucleosome dyad (equivalent to the positional error function in our model). We ran the published sequence-independent deconvolution tool “single template model” from Brogaard et al. (2012) on the Watson and Crick strand fragment ends separately, i.e. distinguishing which side of the cut site the fragment lies on. The nucleotide-frequency profiles surrounding the two dyad datasets reveal an asymmetric preference that cannot be true due to the nucleosome’s point symmetry around the dyad position (Figure 5). Because of the nucleosome’s symmetry the reverse complement sequence has the same binding energy and the asymmetry must stem from an experimental bias. While this asymmetry proves the presence of a sequence bias, it does not reveal how the bias is distributed between the sides and to what it is positioned.

**Figure 5:**
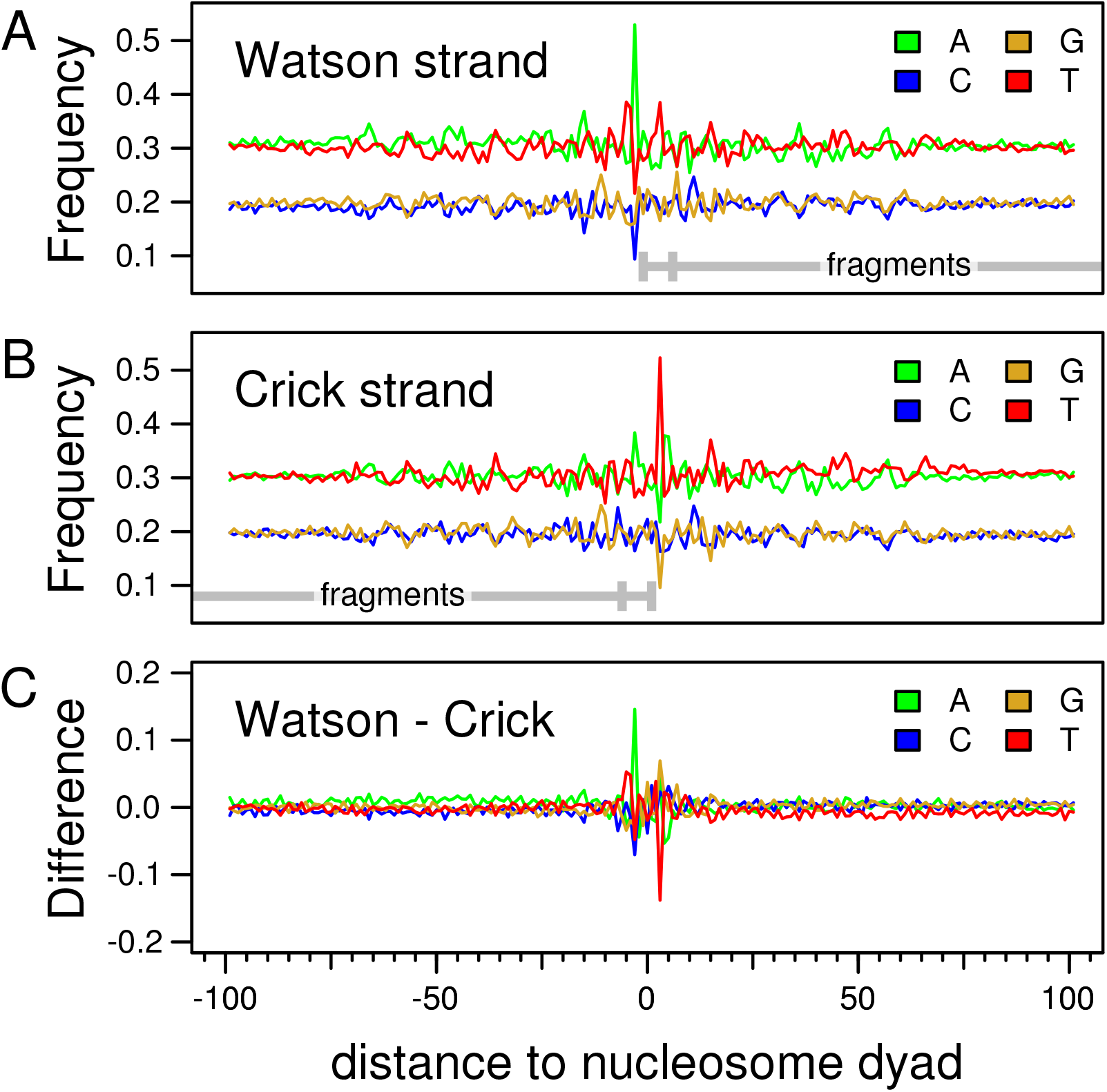
CC-Seq has a strand-specific sequence bias. (**A**, **B**) Nucleotide frequency profiles along the nucleosome region weighted by the Watson and Crick strand CC-Seq measurements. With the tool Brogaard et al. (2012) provide, we separately deconvoluted the reads mapped along the Watson and Crick strand, i.e. the fragment’s start and end in relation to the genome direction (depicted in gray below the subfigures). We weighted the nucleotide occurrences with the computed ‘dyad position’ scores. (**C**) Difference between the frequency profiles: (A) minus (B). The peaks around the nucleosome dyads reveal strong sequence biases close to the cut sites. There is also a minor general depletion of T over the region of the sequenced fragment.

To separate out the bias near the dyad from the real binding preference, we extended our method with two modules that model the sequence bias: positioned in relation to measured nucleosome dyads, or independent measurements unrelated to nucleosomes (see Supplemental Methods). The sequence bias is modelled equivalent to the nucleosome binding energy, but is added in the model of the measurements instead of the thermodynamic model. The module that depends on the dyad position separates the bias from the nucleosome binding preference. Therefore, the method learns a cleaner energy model and shows that the enrichment of A and T in CC-Seq and not its absence elsewhere is the bias (Figure 6A, B). However, the correlation between the extended method and CC-Seq data is not significantly improved.

**Figure 6:**
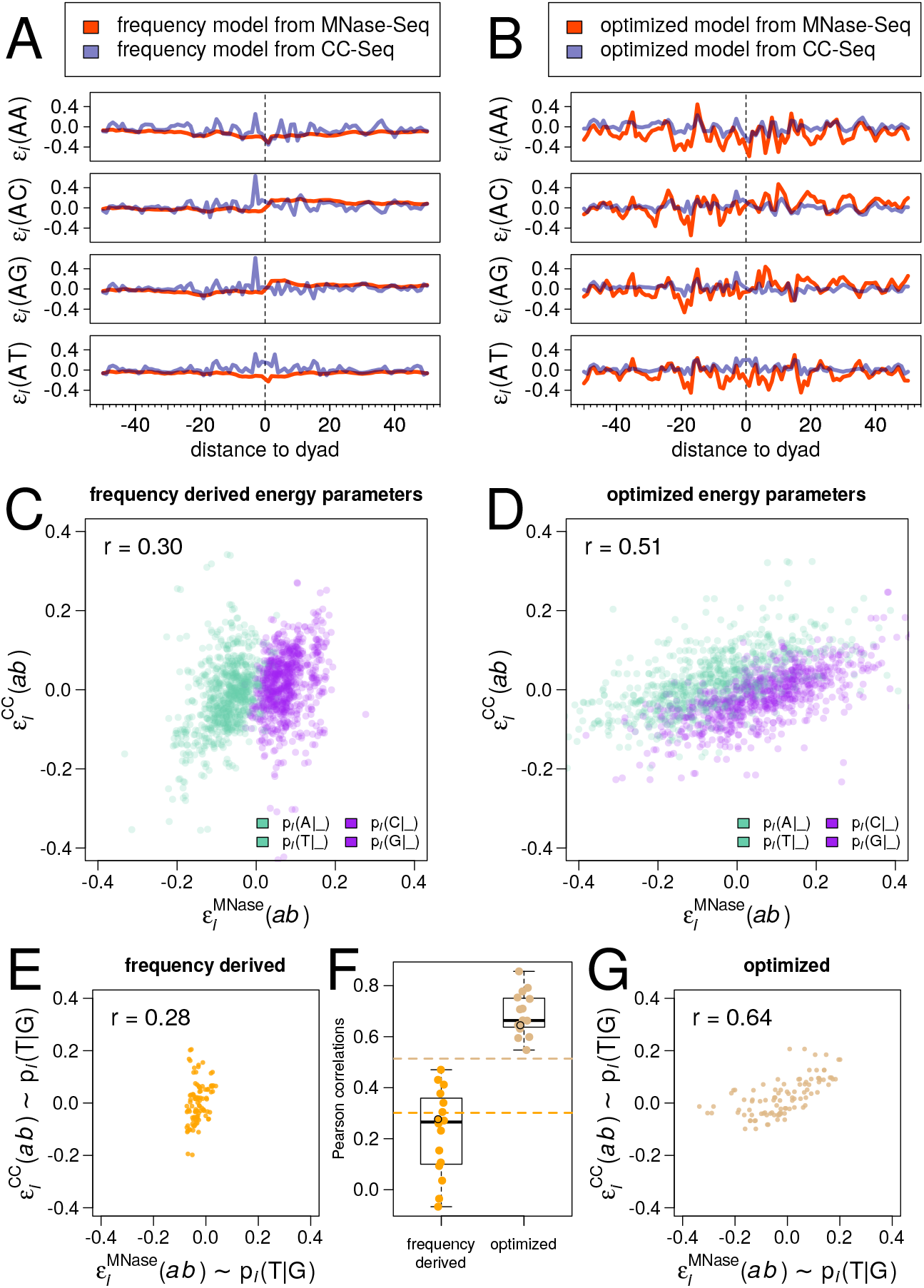
The energy models optimized with our method using MNase-Seq or CC-Seq data are more similar than frequency-derived models. Comparison of the energy models obtained directly from dinucleotide frequencies (**A**, **C**) and by our method (**B**, **D**) using MNase-Seq or CC-Seq data. (A, B) Profiles of a subset of the energy parameters *ε*_*i*_(*ab*) over the nucleosome region. (C, D) Scatterplot of all energy parameters obtained from the two datasets with coloring based on if they represent a conditional probability of G/C or A/T, to visualize the different preferences of G+C content. (**F**) Correlations between the MNase-Seq and CC-Seq based models. Each dot shows the correlation of energy parameters that represent one dinucleotide conditional probability over all positions and the dashed lines show the correlations of all energy parameters. Orange on the left compares the models derived directly from the frequencies and brown on the right compares the optimized models. (**E**, **G**) Scatterplots of the energy parameter comparison marked by black circles in (F).

Because CC-Seq measures nucleosome positioning by cleaving the DNA close to the dyad, a dependence of the cleavage efficiency on the local sequence is reasonable. The cut frequency and the cut-site distribution affect the deconvolution score of a nucleosome. Brogaard et al. (2012) observed that the distribution between the preferred −1 and +6 cut sites depends on the occurrence or absence of A at the −3 and T at the +3 positions and therefore used four separate cut-site distributions to deconvolve the data with the intention of reducing possible biases. While repeating the analysis with this “four template model” improved the similarity between the strand data, the A/T bias is still present (Supplemental Figure S3). Their sequence-dependent deconvolution apparently equalizes the bias of either strand instead of removing it.

### 4.5 Comparisons of energy models obtained from MNase-Seq and CC-Seq

We initialize the dinucleotide binding energy contributions *ε*_*i*_(*ab*) from the dinculeotide frequencies at nucleosome binding sites measured by MNase-Seq or CC-Seq. The energy parameters represent conditional probabilities of one nucleotide given the other (see Methods for the exact definition). The resulting frequency derived parameters differ strongly between the models of MNase-Seq and CC-Seq (Figure 6A, C). The amplitudes of the MNase-Seq model look like a smoothed version of the CC-Seq model, which is expected given the lower resolution of MNase-Seq measurements. The energy model of CC-Seq contains the biased A enrichment at −3, which is absent in the MNase-Seq model. This leads to a low Pearson’s correlation coefficients of 0.30 over all nucleosome binding model parameters *ε*_*i*_(*ab*) between the two models (Figure 6C). The correlation coefficients between parameters *ε*_*i*_(*ab*) of individual conditional probabilities along the model positions span a low range between *−*0.07 and 0.47 (Figure 6F).

In contrast, after training our models the energy parameters obtained from the two datasets are more similar (Figure 6B, D). The amplitude of the MNase-Seq energy model increases more than 4-fold with position-specific features. This reveals that the commonly described 10-bp-periodic pattern is a smoothed version of the position-specific nucleosome preference due to the low-positional resolution of MNase-Seq. In the CC-Seq model the enrichment of A at the −3 positions is separated out of the nucleosome-energy parameters and described with the sequence-bias parameters. This further confirms that the enrichment is an experimental bias. The correlation coefficient between the two models improves from 0.30 to 0.51 and the correlations for conditional probabilities of individual nucleotide pairs improve even more, increasing into a range of 0.54 to 0.86 (Figure 6F).

While the two energy models learned by our method agree much better than the frequency derived models, there are still systematic differences between them (Figure 6B, D). One of the two main differences is the tendency of the MNase-Seq model to have higher *ε*_*i*_(*ab*) values for conditional probabilities of G or C (i.e. p_*i*_(G|_) or p_*i*_(C|_)), while the CC-Seq model has the opposite tendency (Figure 6D). The different preferences of G+C vs A+T content reflect the datasets’ different correlations to G+C content (Figure 2A, purple box). We cannot learn such a general G+C bias well without independent information, because such a bias is hard to distinguish from the nucleosome binding preferences. The second main difference between the models is the roughly 3-fold larger amplitude of the MNase-Seq model’s energy parameters (Figure 6B, D). This larger amplitude might reflect stronger apparent nucleosome binding preferences *in vitro* than *in vivo* due to the additional active processes *in vivo*.

## 5 Discussion

### 5.1 The average genomic nucleosome occupancy has not been measured experimentally

The nucleosome occupancy at a genomic position is the fraction of cells in which the position is covered by a nucleosome (Struhl and Segal 2013). We showed that occupancies cannot be directly derived from genome-wide nucleosome measurements (Figure 3). In fact, the average genomic nucleosome occupancy is still unknown, as all values mentioned in literature are only approximations. A common estimate is that 80% of the genome is covered by nucleosomes (Lee et al. 2007; Shivaswamy et al. 2008). However, this coverage value is based on a limited set of called nucleosomes that are assumed to always be present, whereas in reality nucleosome binding is not binary over the whole population and most genome region are only occupied sometimes.

An average genomic nucleosome occupancy of 75-90% has been given (Field et al. 2008; Segal and Widom 2009; Locke et al. 2010; Liu et al. 2014; Chereji and Morozov 2014), citing the *Chromatin* book of 1989 by van Holde (Holde 1989). However, the book never mentions such a range nor an experiment that measured the average genomic nucleosome occupancy at all. The 75-90% range first appears as an approximation given by Segal et al. (2006), who cite Holde (1989) for *in vitro* measurements their approximation relies on – not a direct measurement. Their approximation assumes that the complete genome is covered by nucleosomes spaced by linkers, which have a length of 10-50 bps based on *in vitro* nucleosome array formation (Holde 1989). To improve this approximation we extended the back-of-an-envelope calculation to include nucleosome-depleted regions, nucleosome binding frequencies, and *in vivo* linker lengths (Supplemental Information 1.1). We estimate an average genomic nucleosome occupancy of 68-83%. Knowing the average nucleosome occupancy is of fundamental importance, and an experiment to measure it directly is long overdue. As we showed the average occupancy limits the possible occupancy distributions. The distribution informs the nucleosome binding energies and, therefore, we cannot learn precise sequence preferences of nucleosomes without knowing the average occupancy.

### 5.2 Do nucleosomes really prefer G+C-rich DNA?

It is generally agreed that G+C-rich DNA is preferred by nucleosomes, but to what degree is still unclear (Iyer 2012; Struhl and Segal 2013). *In vivo* nucleosome measurements in *S.cerevisia* disagree on their correlation to local G+C content (Figure 2). Many datasets show a G+C enrichment in nucleosome-covered sequences, which suggests a strong preference, but at least for MNase-Seq datasets the signal may partially come from sequence biases, because nucleosome-free MNase-Seq measurements show similar G+C enrichments (Figure 2) (Locke et al. 2010; Chung et al. 2010). Similarly, the high correlations between *in vitro* and *in vivo* MNase-Seq datasets suggest that the DNA sequence is important for nucleosome positioning (Kaplan et al. 2009), but the sequence bias of MNase-Seq could contribute to the correlation (Locke et al. 2010). An enrichment of G+C-rich sequences is also seen in *in vitro* measurements of competition between synthetic oligonucleotides to bind histones (Kaplan et al. 2009; Levo et al. 2015). These measurements may have G+C biases of their own, e.g. from the salt-gradient dialysis (Chung et al. 2010).

While the majority of measurements suggest that G+C content influences nucleosome occupancy *in vivo*, several other *in vivo* measurements show low or negative correlations between nucleosome occupancy and G+C content. Three of the *S.cerevisia* datasets we analyzed have low or negative correlation coefficients with G+C content (Shivaswamy et al.: 0.02, Cole et al.: −0.08, Fan et al.: 0.06; Figure 2). CC-Seq measurements in *S.pombe* anti-correlate with G+C content (Moyle-Heyrman et al. 2013). However, these weak correlations with G+C content could also be effects of experimental sequence biases.

While the *S.cerevisia* CC-Seq dataset has a correlation coefficient of 0.35 to G+C content, the predicted occupancies of our optimized CC-Seq model have a strongly negative correlation coefficient of −0.47 to G+C content. The preference of the CC-Seq model for A+T-rich sequences is also seen in the model’s parameters (Figure 6). This probably originates from the thermodynamic model capping the maximal possible occupancy. The method is forced to learn an energy model that is positionally precise instead of replicating the unrealistic large-scale variations that correlate with G+C content. Comparing the model’s predictions with the CC-Seq dataset confirm this: the correlation coefficient is 0.37 for dyad positioning (single-base-pair resolution) and 0.04 for occupancies (147-bp resolution). That our CC-Seq model prefers A+T content, even though the data has a G+C enrichment is an indication that the nucleosomes do not strongly prefer high G+C content in *S.cerevisia*.

Nucleosomes might not prefer G+C-rich sequences even if the nucleosome occupancy correlates with G+C content. The G+C content of nucleosomal DNA was shown to enrich over evolutionary time, caused by nucleotide mutation rates that differ between nucleosome-bound and -unbound DNA (Chen et al. 2012). Support for this hypothesis is a low correlation of G+C content to *in vitro* MNase-Seq measurements of nucleosomes assembled on the genome of *E.coli*, which natively has no nucleosomes (Xing and He 2015). The G+C content also acts as a proxy for other correlated, biologically-relevant sequence features like the physical properties of DNA, which depends primarily on its dinucleotide-composition (Tillo and Hughes 2009). Taken together, the preference of nucleosomes for G+C-rich sequences is unlikely to be the cause for high correlations between nucleosome measurements and G+C content.

### 5.3 Position-specific nucleosome binding preferences

The nucleosomes rotational positioning is influenced by a 10-bp-periodic pattern of WW/SS dinucleotide enrichment (Struhl and Segal 2013). A more jagged pattern was observed with CC-Seq (Brogaard et al. 2012). We showed that, except for the biased peaks at the positions ±3 bps from the dyad, this jagged pattern is closer to the truth than the 10-bp-periodic pattern. The high-resolution energy model our method learned from MNase-Seq data shows a similarly jagged pattern (Figure 6), revealing that the smooth 10-bp-periodic pattern is due to the low resolution of MNase-Seq. The jagged pattern shows the 10-bp periodicity, but the deviations are not periodic themselves.

The low-resolution energy models derived by others (Kaplan et al. 2009; Lubliner and Segal 2009; Xi et al. 2010; Locke et al. 2010), often smoothed on purpose, lead to predictions with lower positional resolution. Our method can learn a high-resolution energy model from low-resolution measurements leading to nucleosome-position predictions with high positional resolution. A nucleosome binding energy model with a higher resolution improves the precision of the thermodynamic model predictions and reduces the errors of the binding energies. Therefore, we hope that our method and similar approaches will reinvigorate quantitative modeling of the competitive binding of nucleosomes and transcription factors at promoters and enhancers to predict gene expression.

### 5.4 How well can we predict nucleosome positioning

Measurements of different experimental protocols disagree on the nucleosome occupancy and binding preferences (Figure 2). The cause for this disagreement are different experimental biases that have effect sizes comparable to those of experimental conditions such as knock-downs of chromosomal proteins or even removal of the whole cellular context (Figure 2). Similar biases in measurements performed with the same experimental protocol or by the same laboratory inflate the correlations between these measurements. Biases shared by the training and test datasets are also responsible for too optimistic results of nucleosome-prediction methods (Kaplan et al. 2009; Tillo and Hughes 2009): we showed that the accuracy can drop dramatically when evaluated on datasets measured with different experimental methods (Supplemental Figure S2).

Low-resolution predictions can outperform high-resolution predictions when benchmarked against low-resolution measurements even though the high-resolution models are more precise. As a case in point, the correlations usually increase the more predictions and measurements are smoothed. Therefore, we use single-base-pair resolution – without smoothing – to benchmark the prediction methods (Figure 4). Often the predictions and dyad measurements are smoothed over a 147 bp running window, which corresponds to comparing occupancies. This smoothing deemphasizes high-precision models and instead emphasizes large-scale feature, which our analysis showed contain strong biases and represent unrealistic genomic occupancy distribution (Figures 2 and 3).

Our two optimized models still disagree in two vital aspects: the amplitude of the pattern and the influence of the average G+C content. To settle this disagreement the method would need to model the experimental bias more precisely. The sequence preference (Fan et al. 2010) and continuous nature (Weiner et al. 2010) of the MNase digestion would need to be modeled for MNase-Seq data, and whole fragments, instead of independently processed fragment ends, would need to be modeled for CC-Seq. A single nucleosome binding energy model could also be learned from distinct experiments simultaneously, optimizing the experimental bias and positional error models for each experiment in parallel.

## 6 Conclusion

We argue that individual high correlations of prediction methods to measurements may feign a better understanding of nucleosome positioning than we have. The methods can be good at predicting specific experimental datasets, but this data is not necessarily a good representation of the biological signal we are interested in. We believe the primary goal of nucleosome-position prediction is to understand what defines the nucleosome positioning *in vivo* in order to create a quantitative model. While experimental biases are widely discussed and their influence is accepted in the field, they have been ignored when formulating quantitative models of nucleosome binding. These fundamental issues still need to be resolved before we can confidently claim to have a good understanding of the sequence preferences of nucleosomes.

We believe that the next major improvements in modeling nucleosome positioning will come from explicitly including experimental errors to separate them from the nucleosome binding energy model. To fully understand what influences nucleosome binding *in vivo* we will have to improve the experimental bias models and integrate extensions of the thermodynamic model into our approach. Additionally, improving the experiments to remove biases or measuring genome-wide absolute nucleosome occupancies would allow us to more critically asses nucleosome-positioning models. We still lack crucial information like the average genomic nucleosome occupancy, which would drastically restrain the nucleosome-position predictions. In any case, integrated models such as ours that combine modelling the biophysics and experimental errors are needed to deal with noise and bias in biological data.

