## Supplemental Methods for "Deconvolving nuclesome binding energy from experimental errors"

### 1 Supplement Methods

For a comprehensive description of the model and analysis see Heron (2017).

#### 1.1 Thermodynamic Model

To describe nucleosome positioning, we use a thermodynamic model similar to those use by others. The model we use incorporates two main aspects of nucleosome positioning: the sequence preference of nucleosomes and the steric exclusion between neighboring nucleosomes. Compared to the thermodynamic models others use, our model describes the probability of a nucleosome dyad occurring at one position compared to other genomic positions (Equation 1). The probabilities differ by their normalization from the commonly used probabilities of a nucleosome dyad occurring at a position or not. Therefore, they are only rescaled by a constant factor (division by the sum over all positions). We had to reformulate the probabilities in this way to define our nuclosome-positioning model.

In our thermodynamic model the probability of a nucleosomes bound at position $i$ is the sum of statistical weights of all configuration that have a nucleosome bound at position $i$ ($Z_i^*$) normalized that $\sum_{i=1}^{L_S} Z_i^* = 1$ with $L_S$ the length of the sequence $S$. In all equations the nucleosome positions will be represented by their dyad position.

$$
\begin{aligned}
\text{P(nucleosome at i}|S,\varepsilon,\mu) &= \frac{Z_i^*}{\sum_{i'=1}^{L_S} Z_{i'}^*} \\
Z_i^* &= F_i^* B_i^* \\
\text{P(nucleosome at i}|S,\varepsilon,\mu) &= \frac{F_i^* B_i^*}{\sum_{i'=1}^{L_S} F_{i'}^* B_{i'}^*}
\end{aligned} \tag{1}
$$

Where $\varepsilon$ and $\mu$ are nucleosome binding energy parameters.

Computing the statistical weight of all legal configurations individually is impractical, because the time complexity is exponential. The Forward/Backward algorithm – a dynamic programming method – reduces the time complexity down to linearity.

For the thermodynamic model, every $Z_i^*$ depends on all other $Z_i^*$. $Z_i^*$ influences the frequency of legal configurations with and without a nucleosome at position $i$, which in turn affects the neighboring statistical weight and so on. Given the linear nature of the DNA and assuming there are no nucleosome interactions beyond the neighbors, the probability scores $Z_i^*$ can be split into two parts that rely on the nucleosome before or after the position $i$. The two parts are called forward ($F_i^*$) and backward ($B_i^*$), because they can be computed by going over the genome linearly forwards and backwards, respectively.

### 1.2 Forward/Backward Method

The Forward ($F_i^*$) and Backward ($B_i^*$) terms are symmetric, with the exception that $F_i^*$ contains the nucleosome binding energy for position $i$, while $B_i^*$ does not. The definitions vary between publications: others have used completely symmetrical Forward and Backward terms by splitting out the binding energy of position $i$ into a third term (e.g. Field et al. (2008)). We provide all equations for completeness, but will only explain the Forward equations for brevity.

$F_i^*$ is the sum of statistical weights of all legal configurations from the left ($< i$) direction. To compute $F_i^*$ only the information of positions in that direction ($< i$) is needed. The equations also use $F_i$, the forward sum of statistical weights for all configurations with no nucleosome covering position $i$ (i.e. it being open).

$$
\begin{aligned}
F_0 &= 1, \quad F_0^* = 0 \quad \text{(Initialization)} \\
F_{i+1} &= F_i + F_{i-D_N}^* \\
F_{i+1}^* &= F_{i-D_N} e^{E_{i+1}-\mu} \\
\\
B_{L_C+1} &= 1, \quad B_{L_C+1}^* = 0 \quad \text{(Initialization)} \\
B_{i-1} &= B_i + B_{i+D_N}^* e^{E_{i+D_N}-\mu} \\
B_{i-1}^* &= B_{i+D_N}
\end{aligned}
\tag{2}
$$

Where $D_N$ is the half length of nucleosome covered DNA (i.e. 73 bps), $E_i$ is the sequence-specific binding energy of a nucleosome at position $i$ and $\mu$ is the sequence-unspecific binding energy of a nucleosome. A position can be open ($F_{i+1}$), either when the previous position is open ($F_i$) or when a nucleosome ended on the previous position ($F_{i-D_N}^*$). For a nucleosome dyad to occur ($F_{i+1}^*$) the position half a nucleosome before has to be open $F_{i-D_N}$. The statistical weight of the position half a nucleosome earlier being open implies that the positions in between are also open.

#### 1.2.1 Sequence-specific binding energy

The common way to describe the sequence binding preference of transcription factors and other DNA-binding factors are position weight matrices (PWMs) (Stormo 2013). A PWM represents the nucleotide preference at each position assuming independence between all positions. Such PWMs are equivalent to 0th-order Markov chains, which in turn are a type of Markov models (the commonly used term). A higher-order Markov chain loosens the independence assumption by using probabilities that depend on the previous positions. If not specified otherwise, we represent the sequence-specific binding energy with a Markov chain of 1st-order, i.e. the probabilities depend on one previous position. We implemented the method so it can handle higher-order Markov chains and have optimized Markov chains up to 4th-order. To represent the dyad symmetric nature of nucleosome binding the method uses conditional probabilities that depend on the positions towards the center. The energy terms $\varepsilon$ that encode the Markov chain represent the logarithm of the conditional probabilities.

$$E_i := \sum_{j=-D_M}^{D_M} \varepsilon_j(s_{i+j-k}..s_{i+j})$$
$$\varepsilon_j(s_{i+j-k}..s_{i+j}) := \begin{cases} \ln \frac{p_j(s_{i+j}|s_{i+j-k}..s_{i+j-1})}{p^{\mathrm{bg}}(s_{i+j}|s_{i+j-k}..s_{i+j-1})} & \text{if } j > 0 \\ \ln \frac{p_j(s_{i+j-k}..s_{i+j})}{p^{\mathrm{bg}}(s_{i+j-k}..s_{i+j})} & \text{if } j = 0 \\ \ln \frac{p_j(s_{i+j-k}|s_{i+j-k+1}..s_{i+j})}{p^{\mathrm{bg}}(s_{i+j-k}|s_{i+j-k+1}..s_{i+j})} & \text{if } j < 0 \end{cases} \tag{3}$$

$k$ is the order of the Markov chain and $D_M$ is the half size of the energy model, which is generally $\leq D_N$. The method initializes the parameters $\varepsilon$ based on Equation 3 by estimating the probabilities $p$ and genomic background probabilities $p^{\mathrm{bg}}$ from nucleotide frequencies, based on Bolzmann's law. The case separation conserves the dyad symmetry of the energy-model parameters (the cases are shown for a 1st-order model). During optimization we remove systematic shifts of the $\varepsilon$ that represent information contained in neighboring probabilities. This conserves the energy model's representation as a Markov chain.

##### 1.2.2 Sequence-unspecific binding energy

In the model $\mu$ represents the sequence-unspecific binding energy. It contains the general binding preference or chemical potential of a nucleosome, and the nucleosome concentration. Because the two cannot be separated without measuring either in an independent experiment, the model represents both with a single parameter.

#### 1.3 Occupancy

The occupancy is unnecessary to compute the likelihood and optimize the parameters. However, a common validation is to compare predicted and measured occupancies. The occupancy is computed by summing the statistical weights of all configuration where a position is covered by a nucleosome and then dividing it by the sum off all possible configurations. There are two ways to normalize by the sum off all possible configurations:

$$\begin{aligned} \mathrm{Occ}(i) &= \frac{\sum_{j=i-D_N}^{i+D_N} Z_j^*}{\sum_{j=i-D_N}^{i+D_N} Z_j^* + F_i B_i} \\ &\text{or} \\ \mathrm{Occ}(i) &= \frac{\sum_{j=i-D_N}^{i+D_N} Z_j^*}{F_0 B_0} \\ &\propto \sum_{j=i-D_N}^{i+D_N} Z_j^* \\ &\propto \sum_{j=i-D_N}^{i+D_N} \mathrm{P}(\text{nucleosome at } j|S,\varepsilon,\mu) \end{aligned} \tag{4}$$

We used unnormalized, i.e. scale-free, occupancies based on the proportional equations, i.e. smoothed dyad probabilities. The nucleosome-positioning measurements have no absolute scale, therefore the validation has to mask scaling – including the normalization – anyway. We also used "occupancy" prediction computed from $\mathrm{P}(x_n \text{ measured}|S, \varepsilon, \mu, \theta)$, which contains experimental errors. This makes sense when comparing the predictions with data measured by the same or similar experiments as the training data.

#### 1.4 Likelihood

To optimize the model parameters we describe observing the experimental data with a likelihood.

$$\mathcal{L} = \prod_{n=1}^{N} \mathrm{P}(x_n \text{ measured}|S, \varepsilon, \mu, \theta)^{w_n} \tag{5}$$

Where $n$ is the index of the measurements, $x_n$ the position, and $w_n$ the summed weight (e.g. counts) of the measurements at position $x_n$. $S$ is the sequence (e.g. genome), $\varepsilon$ the sequence-specific binding energies, $\mu$ the sequence-unspecific binding energy and $\theta$ all other parameters.

The common implicit assumption that the data directly reflects nucleosome positioning corresponds to setting $\mathrm{P}(x_n \text{ measured}|...) := \mathrm{P}(\text{nucleosome at } i|...)$. This assumption is an oversimplification, which misses experimental biases and uncertainties. For example, the read positions of MNase-Seq are imprecise and only estimate the exact nucleosome position. To describe this positional uncertainty the probability of a measurement at position $x_n$ given a nucleosome at $i$ is separated from a nucleosome occurring at position $i$:

$$\mathcal{L} = \prod_{n=1}^{N} \left[ \sum_{i=1}^{N} \mathrm{P}(x_n \text{ measured}|\text{nucleosome at } i, \theta)\, \mathrm{P}(\text{nucleosome at } i|S, \varepsilon, \mu) \right]^{w_n} \tag{6}$$

The computation of $\mathrm{P}(\text{nucleosome at } i|S, \varepsilon, \mu)$ is part of the thermodynamic model. $\mathrm{P}(x_n \text{ measured}|\text{nucleosome at i}, \theta)$ can represent different experimental biases and positional errors. Here, we explain the concept based on the positional-uncertainty model we used.

The positional-uncertainty point spread function is based on the Laplace distribution. The maximal allowed error region is limited to $\pm\delta = \pm 20$, i.e. the model assumes that a measurement cannot be further than 20bps away from the nucleosome dyad.

$$\mathrm{P}(x_n \text{ measured}|\text{nucleosome at } i, \theta) = \mathrm{P}_{\text{dist}}(x_n - i|\eta) = A_\eta e^{\frac{-|x_n - i|}{\eta}} \tag{7}$$

The parameter $\eta$ is equivalent to the standard deviation of a Laplace distribution and controls the width of the point-spread function. $A_\eta$ is the normalization constant that ensures the probabilities sum to one:

$$\begin{aligned} \frac{1}{A_\eta} &= \sum_{k=x_n-\delta}^{x_n+\delta} e^{\frac{-|x_n-k|}{\eta}} = \sum_{k=-\delta}^{\delta} (e^{\frac{1}{\eta}})^{-|k|} = \sum_{k=0}^{\delta} (e^{\frac{1}{\eta}})^{-k} + \sum_{k=1}^{\delta} (e^{\frac{1}{\eta}})^{-k} \\ &= \frac{(e^{\frac{1}{\eta}})^{-\delta} - (e^{\frac{1}{\eta}})}{1 - (e^{\frac{1}{\eta}})} + \frac{(e^{\frac{1}{\eta}})^{-\delta} - 1}{1 - (e^{\frac{1}{\eta}})} = \frac{2(e^{\frac{1}{\eta}})^{-\delta} - (e^{\frac{1}{\eta}}) - 1}{1 - (e^{\frac{1}{\eta}})} \end{aligned} \tag{8}$$

Inserting Equation 7 into Equation 6 and taking its logarithm brings the full log-likelihood for the model with Laplace-like positional uncertainty to:

$$
\begin{aligned}
\log \mathcal{L} &= \sum_{n=1}^{N} \log \left[ \sum_{i=x_n-\delta}^{x_n+\delta} A_\eta e^{\frac{-|x_n-i|}{\eta}} \frac{F_i^* B_i^*}{\sum_{i'=1}^{L_S} F_{i'}^* B_{i'}^*} \right] w_n \\
&= \sum_{n=1}^{N} \left( \log \left[ \sum_{i=x_n-\delta}^{x_n+\delta} e^{\frac{-|x_n-i|}{\eta}} F_i^* B_i^* \right] - \log \left[ \sum_{i=1}^{L_S} F_i^* B_i^* \right] + \log A_\eta \right) w_n \qquad (9) \\
&= \sum_{n=1}^{N} \left( \log \left[ \sum_{i=x_n-\delta}^{x_n+\delta} e^{\frac{-|x_n-i|}{\eta}} F_i^* B_i^* \right] w_n \right) - \log \left[ \sum_{i=1}^{L_S} F_i^* B_i^* \right] \sum_{n=1}^{N} w_n + \log A_\eta \sum_{n=1}^{N} w_n
\end{aligned}
$$

### 1.5 Likelihood maximization

The likelihood is maximized with the help of the partial derivatives. Maximizing the likelihood over a whole datasets is to time-consuming, instead we used a mini-batch gradient ascent. Note that to keep the derivatives simpler the log-likelihood use the natural logarithm, i.e. log := ln for all equations.

##### 1.5.0.1 Gradient of log-likelihood

The partial derivative follow by methodical application of the derivative rules to Equation 9.

$$
\frac{\partial \log \mathcal{L}}{\partial \varepsilon, \mu} = \sum_{n=1}^{N} \frac{\sum_{i=x_n-\delta}^{x_n+\delta} e^{\frac{-|x_n-i|}{\eta}} \left( B_i^* \partial F_i^* + F_i^* \partial B_i^* \right)}{\sum_{i=x_n-\delta}^{x_n+\delta} e^{\frac{-|x_n-i|}{\eta}} F_i^* B_i^*} w_n - \frac{\sum_{i=1}^{L_S} \left( B_i^* \partial F_i^* + F_i^* \partial B_i^* \right)}{\sum_{i=1}^{L_S} F_i^* B_i^*} \sum_{n=1}^{N} w_n
$$

$$
\frac{\partial \log \mathcal{L}}{\partial \eta} = \sum_{n=1}^{N} \frac{\sum_{d=1}^{\delta} \frac{d}{\eta^2} e^{\frac{-d}{\eta}} \left( B_{x_n-d}^* F_{x_n-d}^* + B_{x_n+d}^* F_{x_n+d}^* \right)}{\sum_{i=x_n-\delta}^{x_n+\delta} e^{\frac{-|x_n-i|}{\eta}} F_i^* B_i^*} w_n + \frac{\partial}{\partial \eta} \log A_\eta \sum_{n=1}^{N} w_n \qquad (10)
$$

##### 1.5.0.2 Partial derivatives of Forward/Backward terms

For the sequence-specific binding energies $\varepsilon$ and the sequence-unspecific binding energy $\mu$ the partial derivative of the log-likelihood contains those of the Forward and Backward terms. These Forward and Backward derivatives follow by methodical application of the derivative rules to Equation 2. They are also computed with the Forward/Backward algorithm, because their dependency structure is identical to the originals'.

$$
\begin{aligned}
&\frac{\partial F_0}{\partial \varepsilon_l(q)} = 0, \quad \frac{\partial F_0^*}{\partial \varepsilon_l(q)} = 0 \\
&\frac{\partial F_{i+1}}{\partial \varepsilon_l(q)} = \frac{\partial F_i}{\partial \varepsilon_l(q)} + \frac{\partial F_{i-D_N}^*}{\partial \varepsilon_l(q)} \\
&\frac{\partial F_{i+1}^*}{\partial \varepsilon_l(q)} = \left[\frac{\partial F_{i-D_N}}{\partial \varepsilon_l(q)} + F_{i-D_N}\,\mathrm{I}(s_{i+1+l-k}, ..., s_{i+1+l} = q)\right] e^{E_{i+1}-\mu} \\
&\frac{\partial B_{L_S+1}}{\partial \varepsilon_l(q)} = 0, \quad \frac{\partial B_{L_S+1}^*}{\partial \varepsilon_l(q)} = 0 \\
&\frac{\partial B_{i-1}}{\partial \varepsilon_l(q)} = \frac{\partial B_i}{\partial \varepsilon_l(q)} + \left[\frac{\partial B_{i+D_N}^*}{\partial \varepsilon_l(q)} + B_{i+D_N}^*\,\mathrm{I}(s_{i+D_N+l-k}, ..., s_{i+D_N+l} = q)\right] e^{E_{i+D_N}-\mu} \\
&\frac{\partial B_{i-1}^*}{\partial \varepsilon_l(q)} = \frac{\partial B_{i+D_N}}{\partial \varepsilon_l(q)}
\end{aligned}
\tag{11}
$$

Where $\varepsilon_l(q)$ is the binding-energy parameter of position $l$ for the oligonucleotide $q$ of length $k+1$, i.e. $k$ is the order of the Markov chain that describes the energy model.

$$
\begin{aligned}
&-\frac{\partial F_0}{\partial \mu} = 0, \quad -\frac{\partial F_0^*}{\partial \mu} = 0 \\
&-\frac{\partial F_{i+1}}{\partial \mu} = -\frac{\partial F_i}{\partial \mu} - \frac{\partial F_{i-D_N}^*}{\partial \mu} \\
&-\frac{\partial F_{i+1}^*}{\partial \mu} = \left[-\frac{\partial F_{i-D_N}}{\partial \mu} + F_{i-D_N}\right] e^{E_{i+1}-\mu} \\
&-\frac{\partial B_{L_S+1}}{\partial \mu} = 0, \quad -\frac{\partial B_{L_S+1}^*}{\partial \mu} = 0 \\
&-\frac{\partial B_{i-1}}{\partial \mu} = -\frac{\partial B_i}{\partial \mu} + \left[-\frac{\partial B_{i+D_N}^*}{\partial \mu} + B_{i+D_N}^*\right] e^{E_{i+D_N}-\mu} \\
&-\frac{\partial B_{i-1}^*}{\partial \mu} = -\frac{\partial B_{i+D_N}}{\partial \mu}
\end{aligned}
\tag{12}
$$

The Forward/Backward computations are performed in log-space to cope with large-size increases of the Forward and Backward terms. To perform the computations in log-space the partial derivatives for the sequence-unspecific binding affinity $\mu$ have to be negated to make them positive.

##### 1.5.0.3 Partial derivatives of $A_\eta$

The partial derivative of $\eta$ does not contain the derivatives of the Forward and Backward terms, because they are independent of $\eta$. However, it contains the derivative of the logarithmized normalization term $A_\eta$. $\frac{\partial \log A_\eta}{\partial \eta}$ can be derived from the solved or unsolved summation of

Equation 8:

$$\begin{aligned}\frac{\partial \log A_\eta}{\partial \eta} &= \frac{\partial}{\partial \eta} \log \left( \frac{1 - e^{\frac{1}{\eta}}}{2e^{\frac{-\delta}{\eta}} - e^{\frac{1}{\eta}} - 1} \right) \\ &= \frac{1}{\eta^2} \left( \frac{e^{\frac{1}{\eta}}}{1 - e^{\frac{1}{\eta}}} - \frac{2\delta e^{\frac{-\delta}{\eta}} + e^{\frac{1}{\eta}}}{2e^{\frac{-\delta}{\eta}} - e^{\frac{1}{\eta}} - 1} \right) \end{aligned} \tag{13}$$

or:

$$\frac{\partial \log A_\eta}{\partial \eta} = -A_\eta \sum_{k=-\delta}^{\delta} \frac{|k|}{\eta^2} e^{\frac{-|k|}{\eta}}$$

#### 1.5.1 Position-independent positional-uncertainty point-spread function

To model the cut pattern of CC-Seq we developed a point-spread function where the weight of every position is independent of each other. An independent parameter $\zeta_k$ represents the probability $\mathrm{P}(x_n \text{measured}|\text{nucleosome at } i, \theta)$ for each distance $k$ in the allowed uncertainty region ($\pm\delta$ bp) between the nucleosome and measurement:

$$\mathrm{P}(x_n \text{measured}|\text{nucleosome at } i, \theta) = \mathrm{P}_{\text{dist}}(x_n - i|\zeta) \propto e^{\zeta_{x_n - i}} \tag{14}$$

Leading to the normalization term:

$$\begin{aligned}\sum_{k=-\delta}^{+\delta} e^{\zeta_k} &= \frac{1}{A_\zeta} \\ \mathrm{P}_{\text{dist}}(x_n - i|\delta) &= A_\zeta e^{\zeta_{x_n - i}}\end{aligned} \tag{15}$$

Which leads to the derivative of the log-likelihood:

$$\begin{aligned}\frac{\partial \log \mathcal{L}}{\partial \zeta_j} &= \sum_{n=1}^{N} \frac{e^{\zeta_j} B^*_{x_n+j} F^*_{x_n+j}}{\sum_{i=x_n-\delta}^{x_n+\delta} e^{\zeta_{x_n-i}} F^*_i B^*_i} w_n + \frac{\partial}{\partial \zeta_j} \log A_\zeta \sum_{n=1}^{N} w_n \\ & \mathit{with:} \\ \frac{\partial}{\partial \zeta_j} \log A_\zeta &= \frac{\partial}{\partial \zeta_j} \log \left( \frac{1}{\sum_{k=-\delta}^{+\delta} e^{\zeta_k}} \right) \\ &= \sum_{k=-\delta}^{+\delta} e^{\zeta_k} \frac{-1}{(\sum_{k=-\delta}^{+\delta} e^{\zeta_k})^2} e^{\zeta_j} \\ &= -e^{\zeta_j} A_\zeta\end{aligned} \tag{16}$$

The other terms remain unchanged except for the replaced $\mathrm{P}_{\text{dist}}(x_n - i|\zeta)$.

### 1.6 Sequence-dependent retrieval bias

We developed a model variant that describes a sequence bias to explicitly model the bias we observed in CC-Seq data and separates it from the nucleosome binding model. It uses an energy model (similar to that of a nucleosome) positioned in relation to the dyad position to describe the sequence bias. Given signal described by the sequence bias and nucleosome binding energy overlap, we exploited the difference in symmetry to optimize the two – sequence bias being asymmetric, nucleosome binding energies being dyad symmetric.

$\mathrm{P}(x_n \text{measured}|S, \text{nucleosome at } i, \theta)$ is split into the positional-uncertainty and a sequence-dependent probability to retrieve a measurement from a nucleosome:

$$\begin{aligned}\mathrm{P}(x_n|S,i,\theta) &= \mathrm{P}_{\text{dist}}(x_n - i|\text{retrieved},\theta)\,\mathrm{P}_{\text{seq}}(\text{retrieved}|S,i,\theta)\\ &\quad + \mathrm{P}_{\text{dist}}(x_n - i|\overline{\text{retrieved}},\theta)\,\mathrm{P}_{\text{seq}}(\overline{\text{retrieved}}|S,i,\theta)\\ &= \mathrm{P}_{\text{dist}}(x_n - i|\text{retrieved},\theta)\,\mathrm{P}_{\text{seq}}(\text{retrieved}|S,i,\theta)\end{aligned} \tag{17}$$

The second term drops out, because if no measurement is retrieved than no distance is measured: $\mathrm{P}_{\text{dist}}(x_n - i|\overline{\text{retrieved}},\theta) = 0$. Inserted into the likelihood (Equation 6) we get:

$$\mathcal{L} = \prod_{n=1}^{N}\left[\sum_{i=x_n-\delta}^{x_n+\delta}\mathrm{P}_{\text{dist}}(x_n - i|\text{retrieved},\theta)\,\mathrm{P}_{\text{seq}}(\text{retrieved}|S,i,\theta)\,\mathrm{P}(i|S,\varepsilon,\mu)\right]^{w_n} \tag{18}$$

As with the nucleosomes we use a Markov chain ($C$) to describe the sequence preference that is $\mathrm{P}_{\text{seq}}(\text{retrieved}|i,S,\theta)$. The Markov chain has the weight $\psi_j(s)$ for position $j$ and oligonucleotide $s$:

$$\mathrm{P}_{\text{seq}}(\text{retrieved}|i,S,\theta) = C_{S_i} = e^{\sum_{j=-D_C}^{D_C}\psi_j(s_{i+j})} \tag{19}$$

Where $\pm D_C$ is the extent of the modeled sequence preference. We used $D_C = 10$. Inserted into Equation 18 and logarithmized this leads to:

$$\begin{aligned}\log\mathcal{L} &= \sum_{n=1}^{N}\log\left[\sum_{i=x_n-\delta}^{x_n+\delta}A_\eta e^{\frac{-|x_n-i|}{\eta}}\,\mathrm{P}_{\text{seq}}(\text{retrieved}|i,S)\,\mathrm{P}(i|S,\varepsilon,\mu)\right]w_n\\ &= \sum_{n=1}^{N}\left(\log\left[\sum_{i=x_n-\delta}^{x_n+\delta}e^{\frac{-|x_n-i|}{\eta}}C_{S_i}F_i^*B_i^*\right] - \log\left[\sum_{i=1}^{L_S}C_{S_i}F_i^*B_i^*\right] + \log A_\eta\right)w_n\\ &= \sum_{n=1}^{N}\log\left[\sum_{i=x_n-\delta}^{x_n+\delta}e^{\frac{-|x_n-i|}{\eta}}C_{S_i}F_i^*B_i^*\right]w_n - \log\left[\sum_{i=1}^{L_S}C_{S_i}F_i^*B_i^*\right]\sum_{n=1}^{N}w_n + \log A_\eta\sum_{n=1}^{N}w_n\end{aligned} \tag{20}$$

And the derivative:

$$\begin{aligned}\frac{\partial\log\mathcal{L}}{\partial\psi_{j'(a')}} &= \sum_{n=1}^{N}\frac{\sum_{i=x_n-\delta}^{x_n+\delta}e^{\frac{-|x_n-i|}{\eta}}B_i^*F_i^*\partial C_{S_i}}{\sum_{i=x_n-\delta}^{x_n+\delta}e^{\frac{-|x_n-i|}{\eta}}F_i^*B_i^*C_{S_i}}w_n - \frac{\sum_{i=1}^{L_S}F_i^*B_i^*\partial C_{S_i}}{\sum_{i=1}^{L_S}F_i^*B_i^*C_{S_i}}\sum_{n=1}^{N}w_n\\ \partial C_{S_i} &= \frac{\partial C_{S_i}}{\partial\psi_{j'(a')}} = e^{\sum_{j=-D_C}^{D_C}\psi_j(s_{i+j})}\,\mathrm{I}(S_{i+j'} == a')\\ \frac{\partial\log\mathcal{L}}{\partial\psi_{j'(a')}} &= \sum_{n=1}^{N}\frac{\sum_{i=x_n-\delta}^{x_n+\delta}e^{\frac{-|x_n-i|}{\eta}}B_i^*F_i^*e^{\sum_{j=-D_C}^{D_C}\psi_j(s_{i+j})}\,\mathrm{I}(S_{i+j'} == a')}{\sum_{i=x_n-\delta}^{x_n+\delta}e^{\frac{-|x_n-i|}{\eta}}F_i^*B_i^*e^{\sum_{j=-D_C}^{D_C}\psi_j(s_{i+j})}}w_n\\ &\quad - \frac{\sum_{i=1}^{L_S}F_i^*B_i^*e^{\sum_{j=-D_C}^{D_C}\psi_j(s_{i+j})}\,\mathrm{I}(S_{i+j'} == a')}{\sum_{i=1}^{L_S}F_i^*B_i^*e^{\sum_{j=-D_C}^{D_C}\psi_j(s_{i+j})}}\sum_{n=1}^{N}w_n\end{aligned} \tag{21}$$

#### 1.7 Mini-batch gradient ascent

We used a mini-batch gradient ascent to optimize the model parameters. We defines a batch as all measurements in 25kbp windows, adding 1kbps of sequence information (without measurements) to either side as a buffer. We used a decreasing learning rate and momentum to improve the convergence of the optimization. The optimization is stopped after a maximum of 10k iterations or earlier if the log-likelihood converges/declines.

Because each mini-batch has different data subset, the log-likelihood of one iteration is not comparable with the previous iteration's. Instead of comparing the log-likelihood of recent iterations, we compute the difference to the log-likelihood of the last iteration that trained on the same data subset. By comparing the log-likelihoods after the last and before the new optimization, the computed delta log-likelihood corresponds to the models improvement of describing a subset gained by training on the other subsets. The optimization is interrupted if the average delta log-likelihood of the last 200 iterations drops below a threshold of $-0.01$.

#### 1.8 Runtime

The reasonable computational complexity of the likelihood derivatives together with the application of the Mini-batch gradient ascent allows us to optimize nucleosome binding preference models containing 1.6k parameters in under a day.

#### 1.9 Hyper-parameters:

The default values for the hyper-parameters are listed in a table in the Supplement Information. Due to MNase's cut preference, which leads to biased nucleotide frequencies at the nucleosome borders, we restricted the energy model size to $\pm 50$bps. This excludes the biased nucleosome ends when accounting for the maximal positional uncertainty allowed between the measurements and nucleosomes.
