## Supplemental Information, Figures and Tables for "Deconvolving nuclesome binding energy from experimental errors"

#### 1 Supplement Information

##### 1.1 Estimate of the reasonable average genomic nucleosomes occupancy

We estimated a reasonable average genomic nucleosome occupancy range based on a few basic assumption. The most important assumption is that most of the genome is covered by nucleosomes, which is backed by chromatin accessibility measurements. A similar approximation was used in Segal et al. (2006) to estimate a 75-90% average occupancy. They used linker lengths of 10-50-bps based on nucleosome arrays measurements (Holde 1989). We improved the approximation by using *in vivo* measured linker lengths, including nucleosome-free regions, and allowing for partially absent nucleosomes.

We base the average linker length ($l_{\text{link}}$) of 30 bps on CC-Seq data, which measures dyad to dyad distances directly (Brogaard et al. 2012). We rounded the average linker length upwards, because we are interested in the lower boundary of the nucleosome occupancy. Based on yeast, we use a genome length ($l_{\text{genome}}$) of 12,156,677 bps with 6,275 genes ($n_{\text{genes}}$). As nucleosome length ($l_{\text{nuc}}$) we use the conventional 147 bps.

The average occupancy (occ) of 'fully' covered regions is ~83% and a fully covered genome is theoretically bound by 68682 nucleosomes ($n_{\text{nuc}}$).

$$\begin{aligned} \text{occ} &= \frac{l_{\text{nuc}}}{l_{\text{nuc}} + l_{\text{link}}} = \frac{147}{147 + 30} \approx 0.83 \\ n_{\text{nuc}} &= \frac{l_{\text{genome}}}{l_{\text{nuc}} + l_{\text{link}}} = \frac{12156677}{177} \approx 68682 \end{aligned} \tag{1}$$

Assuming one nucleosome-free region (0% occupancy) per gene gives an average occupancy of ~75%.

$$\text{occ} = \frac{(n_{\text{nuc}} - n_{\text{genes}})l_{\text{nuc}}}{l_{\text{genome}}} = \frac{(68682 - 6275) * 147}{12156677} \approx 0.75 \tag{2}$$

Further assuming that every nucleosome position has a 90% chance of a nucleosome being bound (i.e. 10% absence) gives an average occupancy of ~68%.

$$\text{occ} = \frac{0.9(n_{\text{nuc}} - n_{\text{genes}})l_{\text{nuc}}}{l_{\text{genome}}} = \frac{0.9(68682 - 6275) * 147}{12156677} \approx 0.68 \tag{3}$$

The 90% chance of the average nucleosome to be bound has not been direct measurements for a large sample of nucleosomes. (If that were the case this approximation would probably be unnecessary). MTase measurements at the Pho5 promoter measured frequencies above 90% for three nucleosomes (Small et al. 2014). Based on these measurements we chose the 90% estimate. Chromatin-accessibility assays suggest the genome is mostly covered by nucleosomes and inaccessible, which also suggests high rates of nucleosome binding.

#### 1.2 Additional details for the figures

##### 1.2.1 Figure 2

The Pearson's correlation coefficients are computed between normalized apparent nucleosome occupancies, i.e. derived from the measurements assuming the recovery frequency reflected the nucleosome occurrence frequency. Subfigure A is produced with an adapted version of the corrplot function (Wei 2013).

For the MNase-Seq measurements the fragment centres are approximated from the sequenced fragment ends and used as dyad positions, if no processed dyad positions were published. For the CC-Seq data the published nucleosome scores were used as dyad frequencies. Other datasets from experiments that sequenced the fragments were treated like the MNase-Seq measurements. The dyad frequencies are smoothed over a 147-bp window to approximate nucleosome coverage. For the chip (microarray) experiments, whose measurements already represent coverage, the data was not smoothed. All datasets were normalized to a genome-wide average value of 1.

The preprocessed data is used from the publications and transferred (via the lift-over tool) to the newest yeast genome (sacCer3). The mapping of the raw reads was not repeated to conserve effects different data processing could have. Unmappable genomic regions were excluded from all analysis.

For the G+C content the C+G frequency was smoothed twice with a 147-bp window. The first smoothing produces the G+C content of the nucleosome covered region for each dyad position, and the second smoothing matches the coverage smoothing of the other datasets.

##### 1.2.2 Figure 3

Histograms of the normalized apparent nucleosome occupancies. The datasets were processed as described above for Figure 2.

The Poisson noise distribution was estimated based on the average coverage. This assumes the positions are all independent, which they are not due to the smoothing. However, tests revealed that sampling coverage values directly or smoothing sampled dyads lead to comparable distributions. The approximation is valid and the conclusions hold even if the noise distribution is modelled more precisely.

##### 1.2.3 Figure 4

Pearson's correlation coefficients between dyad probability predictions and dyad position frequencies without smoothing. The dyad position frequencies were derived from the measurements as described for Figure 2 (without smoothing).

The method of Kaplan et al. (2009) was downloaded from their website and ran on the sacCer3 genome with their default parameters.

Our prediction method was trained on the *in vitro* dataset of Kaplan et al. (2009) in the genome direction without symmetrizing the energy model parameters. The predictions where then performed on the reverse complement genome, which is practically independent. To make sure the method was not overfitting, we left one chromosome out of the optimization as an independent validation. The validation on the left-out chromosome showed no signs of overfitting.

###### 1.2.4 Figure 5

The reads mapped to either strand and the deconvolution tool were taken from Brogaard et al. (2012). For this figure the single template model was used to deconvolution the data. For Supplement Figure S3 the four template model was used, which has distinct cut distributions based on the presence and absence of A at the -3 and T at the +3 position in regards to the dyad. The genomic positions are weighted by the computed nucleosome score to compute the nucleotide frequencies, as one would to generate a PWM from numeric measurements. No cutoff or other processing step was applied. The difference values are the frequencies derived from the Crick strand data subtracted from those derived from the Watson strand data.

###### 1.2.5 Figure 6

As described in Supplement Methods 1.2.1, the model parameters $\varepsilon$ are defined in a way to conserve the expected dyad symmetry. For $i < 0$ the conditional probability is from the left nucleotide to the right nucleotide, while for $i > 0$ the conditional probability is mirrored. In subfigures E, F, and G the model parameters are grouped by representing the same conditional probability, i.e. $\mathrm{P}(s_i = a | s_j = b)$ where $j$ is $i + 1$ for $i < 0$ and $i - 1$ for $i > 0$.

##### 1.3 Implementation

All code produced for this publication can be found at https://bitbucket.org/markheron/, https://github.com/markheron/, and/or https://github.com/soedinglab/.

We implemented the method framework in C++ and the time critical parts in C. Time critical parts are maintained in two versions, one that uses normal instructions and one that uses parallel operations on the instruction level (SIMD). This allows for internal validation and speeds up bugfixing. The precision of the normal calculation can be set between single precision (float), double precision (double) and double extended (long double) to check for numerical errors. The SIMD version is only implemented with float SSE2 intrinsics. We further parallelized the partial derivative computations, which are independent of each other, with OpenMP – a multi-threading framework for single computation nodes (OpenMP Architecture Review Board 2008). Thanks to the rise of multi-core CPUs and hundreds of partial derivatives that need to be computed, this can decrease the runtime by an order of magnitude.

We performed the analysis and figure generation in R (R Core Team 2016) with the help of several packages, primarily: Biostrings (Pages et al. 2016), ff (Adler et al. 2014), ffbase (Jonge, Wijffels, and Laan 2015).

#### 2 Supplement Tables

Table S1: Overview of the published nucleosome measurements used in this work. For further details of the experimental protocols we refer to the original publications.

| publication | condition | method | medium | growth | WT strain | cross-linking | measurement |
|---|---|---|---|---|---|---|---|
| Lee et al. (2007) | in vivo | MNase-chip | YPD | log-phase | BY4741 | yes | Affymetrix tiling microarray |
| Shivaswamy et al. (2008) | in vivo / heat shock | MNase-Seq | rich medium | ? | S288C | yes | Solexa sequencing |
| Brogaard et al. (2012) | in vivo | CC-Seq | YPD | log-phase | BY4741 | no | ABI SOLID sequencing |
| Celona et al. (2011) | in vivo / nhp6 KD | MNase-Seq | YPD | ? | S288C | no | Illumina GA IIx |
| Kaplan et al. (2009) | in vivo / in vitro | MNase-Seq | YPD | log-phase | YLC8 | no | Illumina sequencer |
| Gossett and Lieb (2012) | in vivo / H3 KD | MNase-Seq | YPD | log-phase | YEF473A | yes | Illumina GA IIx |
| Zhang et al. (2009) | in vitro | MNase-Seq | - | - | - | no | Illumina GA |
| Cole et al. (2015) | in vivo | MNase-ExoII-Seq | ? | log-phase | YVN381 | no | Illumina sequencer |
| Mavrich et al. (2008) | in vivo | Mnase-ChIP-Seq | rich medium | log phase | BY4741 | yes | 454 pyrosequencing |
| Field et al. (2008) | in vivo | MNase-Seq | ? | log-phase | ? | no | 454 pyrosequencing |
| Fan et al. (2010) | in vivo | ChIP-chip | YPD | log-phase | BY4741 | ? | Affymetrix tiling microarray |
| Locke et al. (2010) | naked DNA | MNase-Seq / Sono-Seq | - | - | - | no | Illumina GA |

Table S2: Optimization parameters

| Parameter | Denomination | Value |
|---|---|---|
| | nr_of_iterations | 10000 |
| | fragment_length | 25000 |
| | fragment_side_buffer | 1000 |
| | numeric_double | 0 |
| | numeric_double_long | 0 |
| | use_SSE | 1 |
| | stopcriterion_deltaLL | -0.01 |
| | stopcriterion_amount | 200 |
| | stopcriterion_min_parameter_change | 0.00 |

Table S3: Learning-rate parameters

| Parameters | Denomination | Value |
|---|---|---|
| $\lambda$ | steplength_general | 0.10 |
| $\lambda^{\varepsilon}$ | steplength_epsilon | 1.00 |
| | steplength_epsilon_lower | 1.00 |
| $\lambda^{\mu}$ | steplength_mu | 15.00 |
| $\lambda^{\eta}$ | steplength_eta | 10.00 |
| | steplength_decay_frequency | 400 |
| $\Lambda$ | steplength_decay_speed | 0.10 |
| $1 - \rho_0$ | exponential_average_old_weight | 0.00 |
| $\varrho$ | exponential_average_increase_speed | 0.50 |

Table S4: Model parameters

| Parameters | Denomination | Value |
|---|---|---|
| $2D_M + 1$ | binders_motif_length | 100 |
| $2D_N + 1$ | binders_footprint | 147 |
| $\mu$ | initial_mu | 4.00 |
| $\delta$ | DELTA | 20 |
| $k + 1$ | mer_length | 2 |

### 3 Supplement Figures

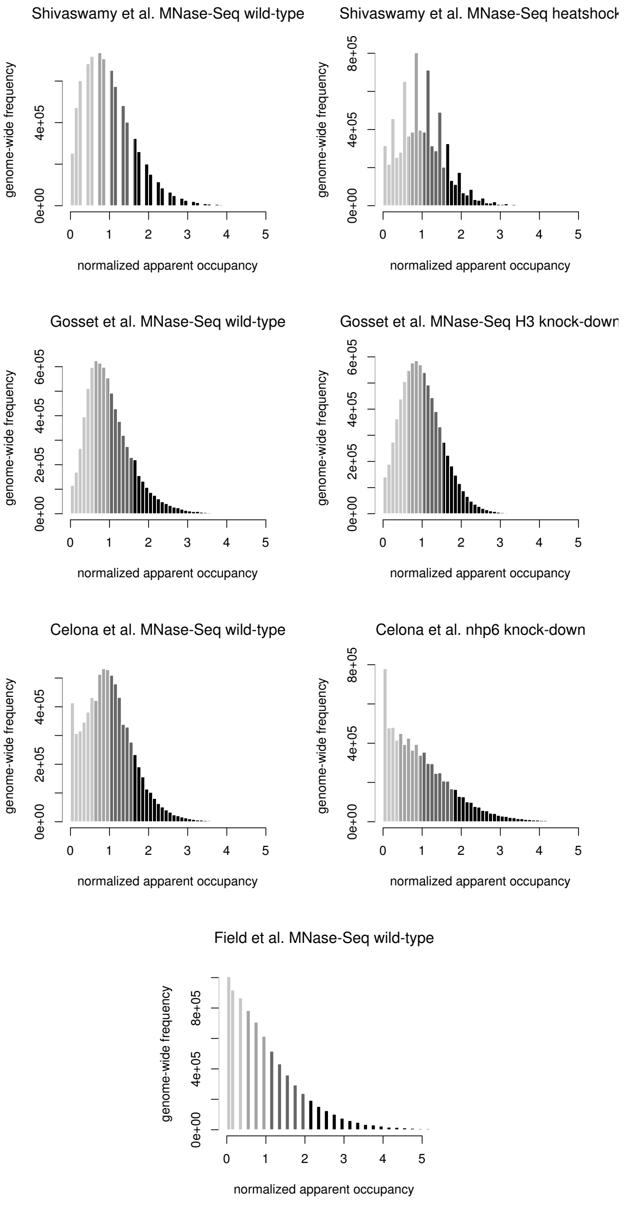


Figure S1: continued on next page

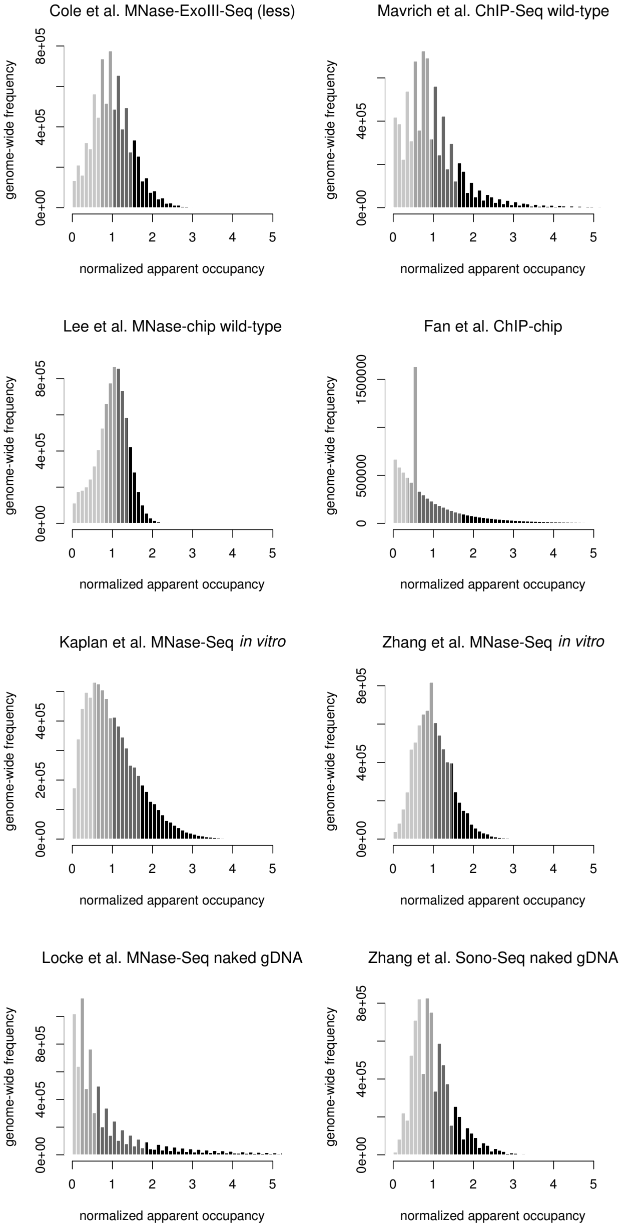


Figure S1: **Nucleosome occupancies distribution of measurements**: Histograms of nucleosome occupancies deduced from measured fragment centers as show in Figure 3A,B for further datasets.

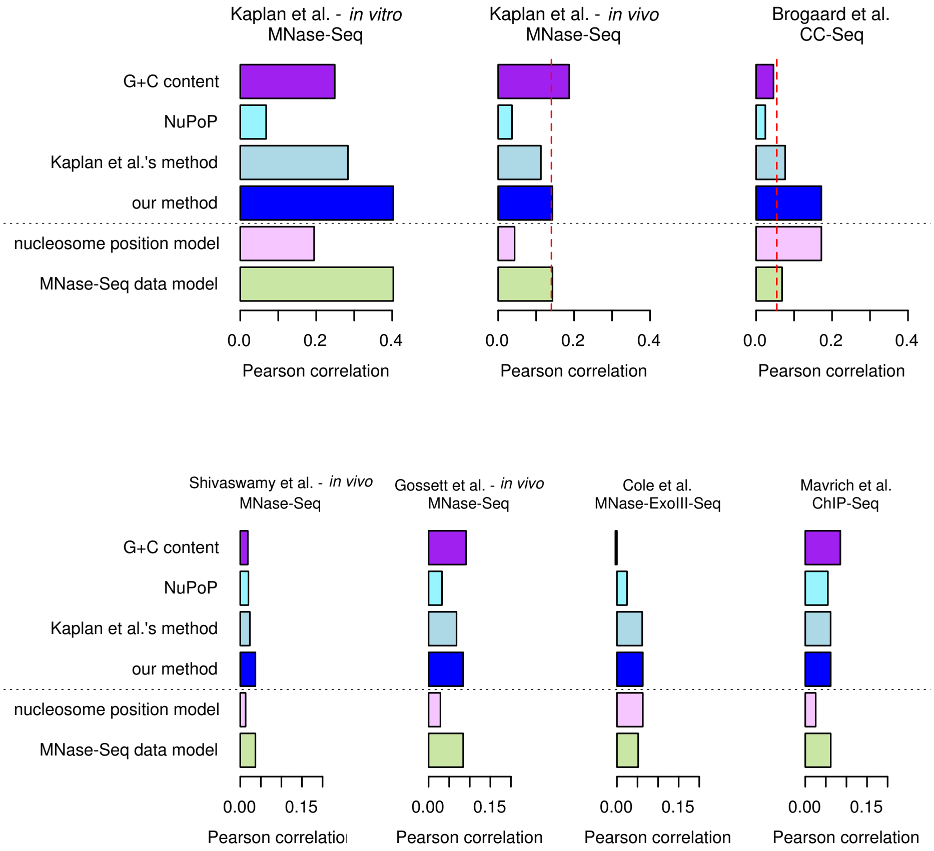


Figure S2: **Further method performances**

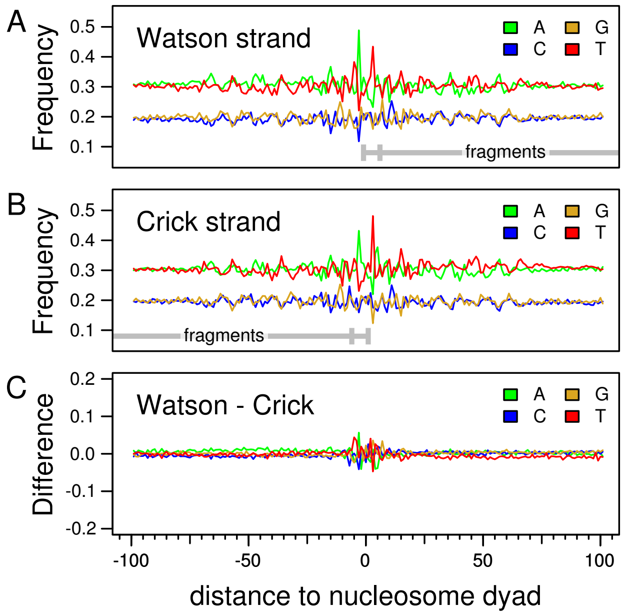


Figure S3: **CC-Seq's sequence bias with the four template model**: Equivalent to **??**, but using the four template model to deconvolute the data insted of the single template model. The difference in (C) is decreased, because the bias in (A) and (B) becomes more symmetrical.

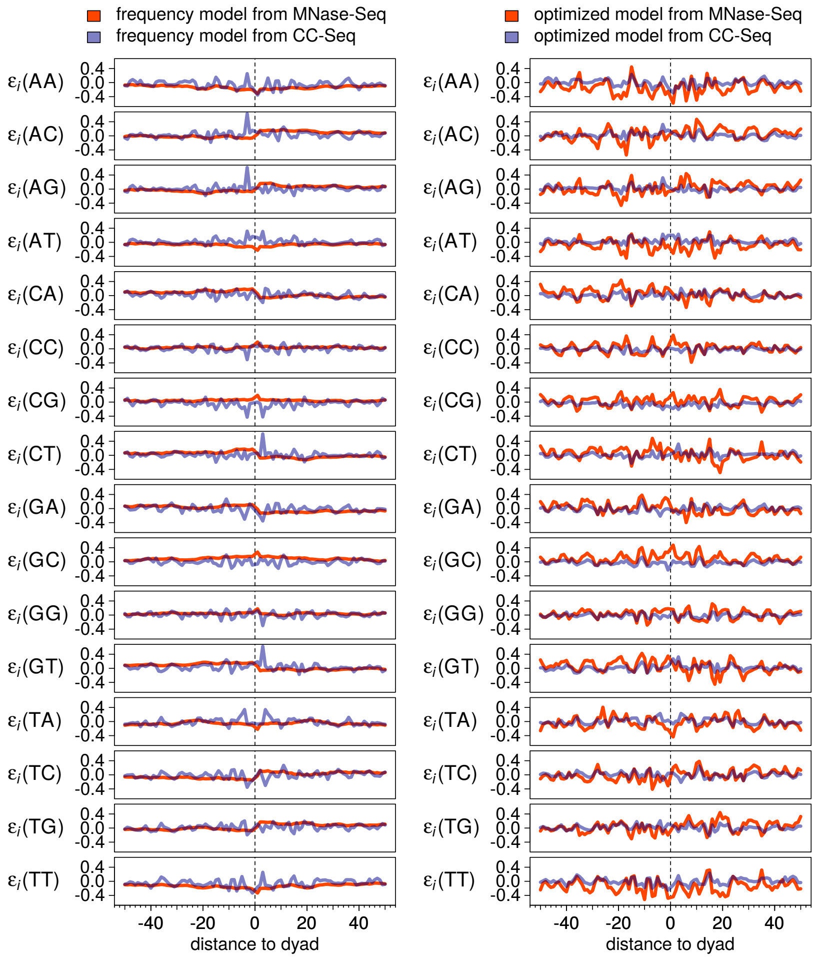


Figure S4: **Full comparison of our energy models.**: All profiles from which a sample is shown in Figure 6A, B.

*Representing Biological Sequences, and Matching Algorithms.*

R Core Team. 2016. *R: A Language and Environment for Statistical Computing.* Vienna, Austria: R Foundation for Statistical Computing. https://www.R-project.org/.

Segal, E., Y. Fondufe-Mittendorf, L. Chen, A. Thåström, Y. Field, I. K. Moore, J.-P. Z. Wang, and J. Widom. 2006. "A Genomic Code for Nucleosome Positioning." *Nature* 442 (7104): 772–78. doi:10.1038/nature04979.

Shivaswamy, S., A. Bhinge, Y. Zhao, S. Jones, M. Hirst, and V. R. Iyer. 2008. "Dynamic Remodeling of Individual Nucleosomes Across a Eukaryotic Genome in Response to Transcriptional Perturbation." *PLoS Biol.* 6 (3): e65. doi:10.1371/journal.pbio.0060065.

Small, E. C., L. Xi, J.-P. Wang, J. Widom, and J. D. Licht. 2014. "Single-Cell Nucleosome Mapping Reveals the Molecular Basis of Gene Expression Heterogeneity." *Proc. Natl. Acad. Sci. USA* 111 (24): E2462–71. doi:10.1073/pnas.1400517111.

Wei, Taiyun. 2013. *Corrplot: Visualization of a Correlation Matrix.* https://CRAN.R-project.org/package=corrplot.

Zhang, Y., Z. Moqtaderi, B. P. Rattner, G. Euskirchen, M. Snyder, J. T. Kadonaga, X. S. Liu, and K. Struhl. 2009. "Intrinsic Histone-Dna Interactions Are Not the Major Determinant of Nucleosome Positions in Vivo." *Nat. Struct. Mol. Biol.* 16 (8): 847–52. doi:10.1038/nsmb.1636.
